# The *Prevotella copri* complex comprises four distinct clades that are underrepresented in Westernised populations

**DOI:** 10.1101/600593

**Authors:** Adrian Tett, Kun D. Huang, Francesco Asnicar, Hannah Fehlner-Peach, Edoardo Pasolli, Nicolai Karcher, Federica Armanini, Paolo Manghi, Kevin Bonham, Moreno Zolfo, Francesca De Filippis, Cara Magnabosco, Richard Bonneau, John Lusingu, John Amuasi, Karl Reinhard, Thomas Rattei, Fredrik Boulund, Lars Engstrand, Albert Zink, Maria Carmen Collado, Dan R. Littman, Daniel Eibach, Danilo Ercolini, Omar Rota-Stabelli, Curtis Huttenhower, Frank Maixner, Nicola Segata

**Author notes:** Correspondence: Adrian Tett; Nicola Segata.

## Abstract

*Prevotella copri* is a common inhabitant of the human gut. Interest in *P. copri* has gathered pace due to conflicting reports on whether it is beneficial or detrimental to health. In a cross-continent meta-analysis exploiting >6,500 available metagenomes supported by new isolate sequencing and recovery of high-quality genomes from metagenomes, we obtained >1,000 *P. copri* genomes. This 100-fold increase over existing isolate genomes allowed the genetic and global population structure of *P. copri* to be explored at an unprecedented depth. We demonstrate *P. copri* is not a monotypic species, but encompasses four distinct clades (>10% inter-clade vs. <4% intra-clade average single nucleotide variants) for which we propose the name *P. copri* complex, comprising clades A, B, C and D. We show the complex is near ubiquitous in non-Westernised populations (95.4% versus 29.6% in Westernised populations), where all four clades are typically co-present within an individual (61.6% of the cases), in contrast to Westernised populations (4.6%). Genomic analysis of the complex reveals substantial and complementary functional diversity, including the potential for utilisation of complex carbohydrates, suggestive that multi-generational dietary modifications may be a driver for the reduced *P. copri* prevalence in Westernised populations. Analysis of ancient stool microbiomes highlights a similar pattern of *P. copri* presence consistent with modern non-Westernised populations, allowing us to estimate the time of clade delineation to pre-date human migratory waves out of Africa. Our analysis reveals *P. copri* to be far more diverse than previously appreciated and this diversity appears to be underrepresented in Western-lifestyle populations.

## Introduction

*Prevotella copri* is a frequently observed inhabitant of the human intestinal microbiome and it displays a large inter-individual variation (Human Microbiome Project Consortium, 2012; Qin et al., 2010; Truong et al., 2017). Being 39.1% prevalent in healthy individuals from current metagenomic profiles (Pasolli et al., 2017), *P. copri* is not ubiquitous. But when present, it is often the most abundant species identified (34% of instances) and it represents the clearest reproducible gut microbiome structure (Arumugam et al., 2011; Koren et al., 2013).

*P. copri* also came to prominence due to its reported association with inflammatory diseases, including new-onset rheumatoid arthritis (Scher et al., 2013), ankylosing spondylitis (Wen et al., 2017) and intestinal inflammation in chronic HIV-1 infection (Dillon et al., 2014). It has also been shown to induce insulin resistance and glucose intolerance (Pedersen et al., 2016). Conversely, others have linked *P. copri* with improved glucose and insulin tolerance in diets rich in fibre (De Vadder et al., 2016; Kovatcheva-Datchary et al., 2015) which suggests the beneficial effects of *P. copri* could be diet dependent (Pedersen et al., 2016). As previously expressed (Cani, 2018; Ley, 2016), such conflicting reports regarding the benefits of *P. copri* suggest that it is an important but enigmatic member of the microbiome.

Higher prevalence of *Prevotella* has been consistently reported in non-Westernised populations (De Filippo et al., 2010; Hansen et al., 2019; Obregon-Tito et al., 2015; Schnorr et al., 2014; Smits et al., 2017; Yatsunenko et al., 2012) and metagenomic studies, capable of species-level resolution, have shown *P. copri* to be particularly prevalent (Pasolli et al., 2019; Vangay et al., 2018). Non-Westernised populations follow a traditional lifestyle and typically consume diets rich in fresh unprocessed food (vegetables and fruits) and high in fibre compared to Western populations (De Filippo et al., 2010; Segata, 2015; Statovci et al., 2017). This association may further support the hypothesis of diet being an important factor in selecting and shaping *Prevotella* populations as it has been shown for the overall diversity of microbial ecosystems (Smits et al., 2017; Sonnenburg and Bäckhed, 2016; Sonnenburg et al., 2016). The high prevalence in societies with more traditional rather than high-fat diets may also support the health benefit of *P. copri*.

Despite the importance of *P. copri* and the open-ended question regarding its role in health and disease there is a lack of available reference genomes and much of what is currently known relies on studies of the single type strain *P. copri* DSM18205 (Hayashi et al., 2007). Recent reports have begun to highlight a degree of strain level heterogeneity within *P. copri* (De Filippis et al., 2019; Truong et al., 2017; Vangay et al., 2018). Indeed, sub-species strain variation may account for at least some of the differences in the reported benefits or detriments of *P. copri*. Yet to date there has been no large scale concerted effort to explore the distribution and genetic variation within *P. copri*.

Here we use a combination of isolate sequencing and large scale metagenomic assembly and strict quality control to recover over 1,000 *P. copri* genomes from publicly available metagenomes spanning multiple host-geographies, disease and lifestyles. We also expand the catalogue of non-Westernised sampled populations with additional metagenomic sequencing of individuals from Ghana, Ethiopia and Tanzania and further profile *P. copri* in ancient intestinal samples from a European natural ice-mummy and stools of pre-Columbian amerinds. These datasets and analyses provide an unprecedented comprehensive insight into the genetic diversity, global population structure and evolutionary history of *P. copri*.

## Results and discussion

### Analysis of >1,000 *P. copri* genomes reveals four clades comprising the *P. copri* complex

To investigate the global distribution and population structures of *P. copri*, we performed an analysis of 6,874 publicly available metagenomes from 36 individual datasets (**Supplementary Table S1**), representing six continents and over 20 different countries. By means of a novel assembly and mapping-based computational approach, we expanded the total number of available *P. copri* genomes to 1,023 and all metagenomically assembled genomes can be defined as high-quality according to current guidelines (Bowers et al., 2017) (estimated completeness >95%, contamination <5%, see **Methods**). This approach, (see **Methods**) involved collating a highly representative set of genomes comprising our newly sequenced *P. copri* isolates (n=15), publicly available references (n=2), as well as a set of carefully curated and manually-guided metagenomically assembled genomes from diverse populations (n=55). This set of 72 genomes was used as a pangenomic reference to bin via mapping metagenomically-assembled contigs from single samples into whole *P. copri* genomes (n=949) (see **Methods**). The obtained genomes were subjected to strict quality control including estimation of within sample strain heterogeneity (see **Methods**) and resulted in genomes with assembly characteristics comparable with those of isolate sequencing and manually curated metagenomic assembled genomes (**Supplementary Figure S1; Supplementary Table S2**). This genome catalogue spans multiple host geographies, populations and lifestyles and can thus be mined to ask fundamental questions regarding the population genomic structure of *P. copri*.

*P. copri* is strikingly revealed by this analysis to be not a monotypic species, but comprised of four distinct clades when placed in phylogenetic context with the closest publicly available representatives of the wider *Prevotella*, *Alloprevotella*, and *Paraprevotella* genera (Figure 1A). These four clades are clearly distinct to other *Prevotella* and to the other considered species, each clade is supported by at least one of our newly sequenced genomes from isolates (Figure 1A) and all clades are represented in our three newly sequenced non-Westernised datasets (98 new genomes, see below). The average nucleotide identity (ANI) distances between the *P. copri* genomes revealed a limited intra-clade distance (mean 2.55% s.d. 0.35% for Clade B to 4.16% s.d. 0.78% for Clade C). Conversely, the inter-clade distances were very high, with values ranging from 13.0% to 21.4%. In comparison, each of the four *P. copri* clades were >23.0% distant to any other *Prevotella*, *Alloprevotella* and *Paraprevotella* species, indicating that the four distinct *P. copri* clades are genetically closer to each other than genomes outside the *P. copri* complex (Figure 1B).

**Figure 1.**
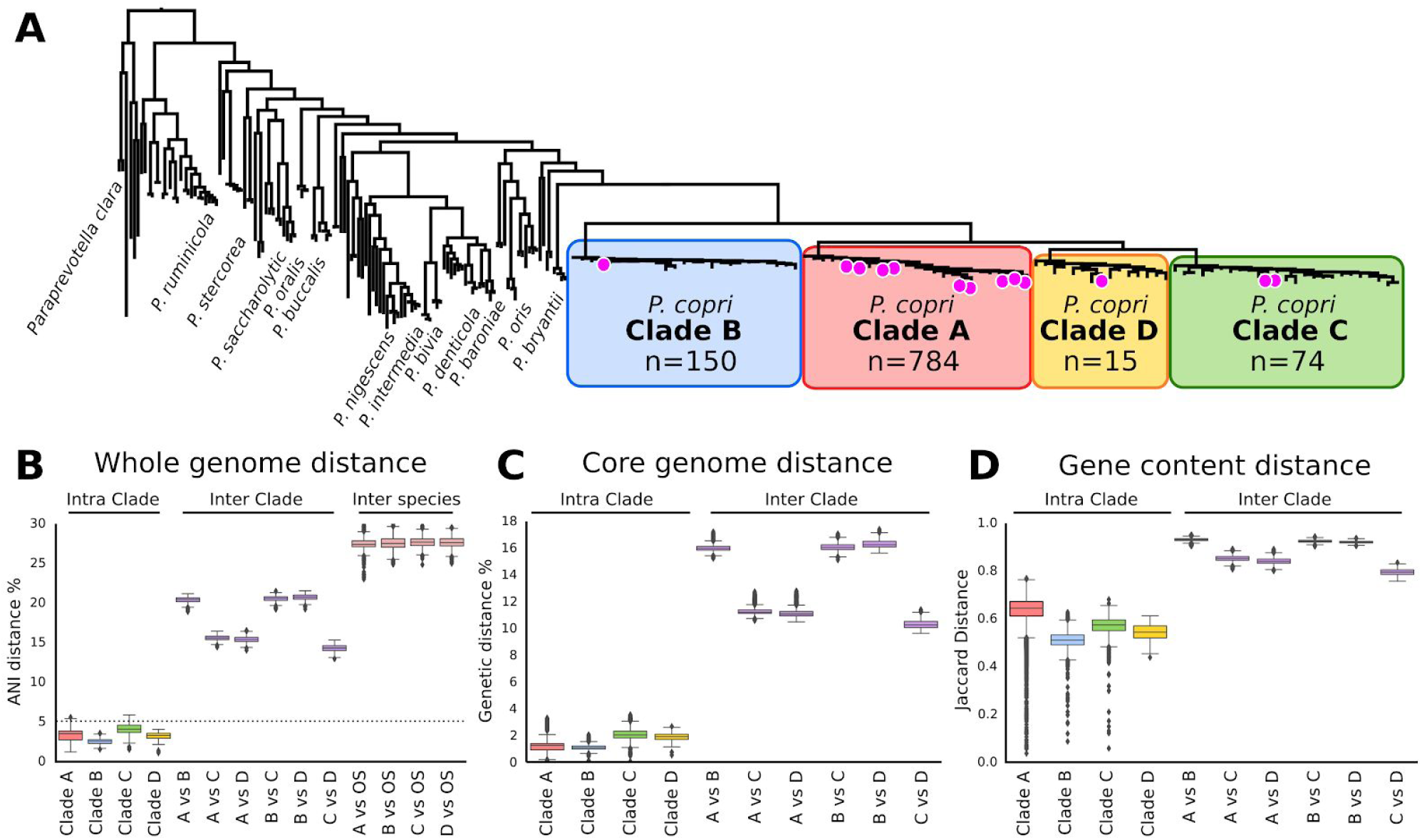
**(A)**. Whole-genome phylogenetic tree of a representative subset of the four distinct *P. copri* clades comprising the *P. copri* complex in relation to other sequenced members of the genera *Prevotella*, *Alloprevotella* and *Paraprevotella*. Magenta circles indicate *P. copri* sequenced isolates. The phylogeny containing all *P. copri* genomes is available as **Supplementary Data File 1** (see Data Availability). **(B**) Genetic distance within clade (intra-clade), between clades (inter-clade) and between clades and other species (denoted OS) of *Prevotella*, *Alloprevotella* and *Paraprevotella* (inter-species), shown as pairwise average nucleotide identity distances (ANI distance). The dotted line denotes 5% ANI distance. **(C-D)** Pairwise SNV distances based on core gene alignment, and Jaccard distance based on pairwise gene content (see **Methods**) between and within the *P. copri* clades.

The high inter-clade genetic distance observed suggests the genome catalogue could represent four distinct species. Studies have sought to place a threshold at which ANI values between genomes equate to the delineation of strains into species, with a broad consensus being values above 5-6% distance (Goris et al., 2007; Jain et al., 2018; Konstantinidis and Tiedje, 2005; Pasolli et al., 2019). All members of all four clades fall well below this threshold when compared to other *P. copri* clades (>10% ANI distance) (Figure 1B). The distinction of the four clades is further supported based on core genome single nucleotide distance (>10% distance) (Figure 1C), by the separation of the clades based purely on gene content (Figure 1D) as well as based on phylogeny (Figure 1A). Respecting the clear distinction of the *P. copri* clades and being conscious of the difficulties in advising separation into species we propose the naming of *P. copri* to encompass these four distinct clades. Therefore we recommend the term “*Prevotella copri* complex” for which there are four genetically distinct Clades A, B, C and D, named sequentially based on the decreasing number of genomes recovered (Figure 1A).

### The four clades are globally distributed with instances of country-specific subtypes

In this study, *P. copri* genomes were recovered from 22 different countries offering a unique opportunity to investigate the biogeographical population structure. These clades were not strictly separated based on geographical location, i.e. all of the four *P. copri* clades were identified in multiple countries and spanning multiple continents (Figure 2). However, within several clades we did observe geographical stratification. In Clade A, for which the most genomes were recovered, we identified three sub-types which were either exclusively or nearly so to samples of Chinese origin, and in addition there was also a cluster exclusive to Israel. In Clade B a specific cluster was identified which can be attributed to Fiji. For Clade C and D it is difficult to ascertain if there is stratification due to the lower number of genomes recovered for these two *P. copri* clades. While geographical stratification was evident for some intra-clade sub-types, most sub-types appeared to be multi-country and even multi-continental, indicating that not only at the clade level but also at intra-clade level *P. copri* is widely geographically distributed.

**Figure 2.**
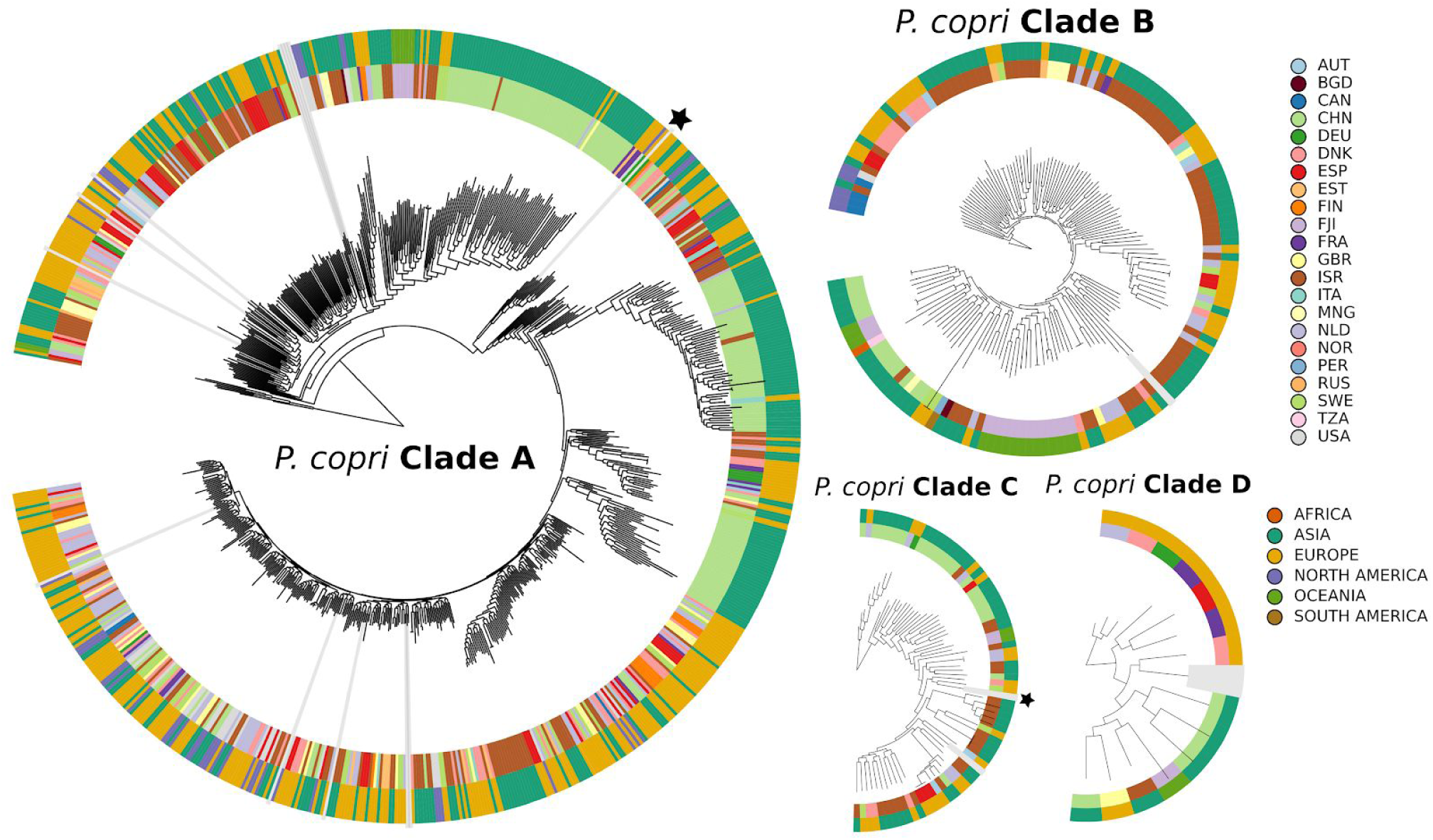
Phylogenetic representation of all 1,023 *P. copri* genomes separated for each clade of the *P. copri* complex. Outer ring is coloured by continent of origin and inner ring is further divided by country. Radial grey bars indicate newly sequenced isolate genomes and publicly available reference genomes are denoted by black stars.

### Associating *P. copri* clades with metagenomically investigated human diseases

A question that remains to be resolved is whether *P. copri* is of benefit or detriment to human health as there are studies reporting both scenarios (De Vadder et al., 2016; Dillon et al., 2014; Kovatcheva-Datchary et al., 2015; Pedersen et al., 2016; Scher et al., 2013; Wen et al., 2017). Here, in a meta-analysis of available disease phenotypes we found no strong indicator that they were associated with the four clades of the complex. Specifically, to investigate the association of the *P. copri* complex with different diseases, we analysed the prevalence and abundance of the four clades for each cohort where the study design included both case and controls. In total there were 10 datasets including colorectal cancer (CRC) (Feng et al., 2015; Vogtmann et al., 2016; Yu et al., 2017; Zeller et al., 2014), type-2-diabetes (T2D) (Karlsson et al., 2013; Qin et al., 2012), hypertension (Li et al., 2017), liver cirrhosis (Qin et al., 2014) and inflammatory bowel disease (IBD) (He et al., 2017; Nielsen et al., 2014).

To identify and estimate the abundance of each of *P. copri* clades within a sample the metagenomic reads were mapped to a panel of unique clade-specific markers designed for each of the four clades (see **Methods**). The most significant changes in abundance and prevalence of *P. copri* and specifically the four clades, were identified in the CRC and adenoma cohort of FengQ et al. with both Clade A and C being associated with disease (**Supplementary Figure S2**). However, three other CRC cohorts considered (Vogtmann et al., 2016; Yu et al., 2017; Zeller et al., 2014) and an overall CRC meta-analysis comprising 7 cohorts failed to support this observation (Thomas et al., 2019). Generally, while there were some weak associations of the *P. copri* clades in disease, across the control samples of the different datasets we observe heterogeneity in both abundance and prevalence suggesting significant batch effects. As such, at the clade level there is no clear evidence to suggest *P. copri* is associated with the etiology of these diseases. We further considered if the distribution of the *P. copri* complex could be associated with other factors such as body mass index (BMI) and age (**Supplementary Figures S3 and S4**). Similar to disease, we note potential batch and cohort effects but no significant differences with all four clades being identified across all age groups and BMI categories.

### Reconstruction of 98 additional genomes from non-Westernised samples expands the diversity of the *P. copri* clades with fewer representatives

Most of our understanding of the microbiome has been accumulated from Westernised populations. While small in comparison to Westernised populations, a number of publicly available datasets have been generated from non-Westernised populations inhabiting Peru (Obregon-Tito et al., 2015), Fiji (Brito et al., 2016), Mongolia (Liu et al., 2016) and two from Tanzania (Rampelli et al., 2015; Smits et al., 2017) totaling 340 metagenomes. The term “Westernisation” encompasses many factors including lifestyle, environment and diet (for full description see **Methods**). A common feature of non-Westernised datasets has been the frequent observation of high *Prevotella* prevalence (De Filippo et al., 2010; Hansen et al., 2019; Obregon-Tito et al., 2015; Schnorr et al., 2014; Smits et al., 2017; Yatsunenko et al., 2012) and particularly *P. copri* in these populations (Pasolli et al., 2019; Vangay et al., 2018). To further investigate the prevalence and abundance of *P. copri* in non-Westernised populations, we also considered our recently sequenced dataset of non-Westernised adults from Madagascar (110 metagenomes) (Pasolli et al., 2019) and three new non-Westernised cohorts. These included paired infant and mother samples from Ethiopia (50 metagenomes) and extended families from Ghana and Tanzania (44 and 68 metagenomes respectively) (see **Methods**). From these additional 272 metagenomes (**Supplementary Table S3**), we recovered 98 high quality *P. copri* complex genomes expanding the clades represented by fewer reconstructed members (Clade A, B, C and D were expanded by 3.4%, 34.7%, 17.6% and 40% respectively) (**Supplementary Figure S5; Supplementary Table S4**). An additional feature of the three newly sequenced datasets was that they included sampling within families. This offered the potential to establish if transmission occurs and if so to what extent within families. When *P. copri* genomes were recovered from more than one family member, we compared the genetic distances to estimate the level of intra-family strain sharing. Using normalised phylogenetic tree distances and cutoffs proposed previously (Truong et al., 2017) (see **Methods**) in 5 of 26 cases (19.2%) we identified the same strain, suggesting both horizontal and vertical transmission of the *P. copri* complex within family units, and therefore familial prevalence of *P. copri* as a potential important source in acquisition.

### Co-presence of multiple *P. copri* complex clades is typical in individuals from non-Westernized populations

We detected the *P. copri* complex in all 40 datasets considered, but the prevalence in non-Westernised populations was nearly ubiquitous (95.4% prevalence) in contrast to Westernised populations (29.6% prevalence) (Figure 3A, **Supplementary Figure S6**). Extending this to consider each of the four *P. copri* clades separately, all four were found to be significantly higher (p-value < 1.1e-12, Welch’s t-test) in the non-Westernised datasets with respect to Westernised, with the *P. copri* complex Clade A being the most prevalent (91.5% in non-Westernized vs 26.9% in Westernised populations) followed by C (88.2% vs 8.35%), B (73.5% vs 6.2%) and D (68.8% vs 2.7%, Figure 3A). The agreement that all four *P. copri* clades are always higher in non-Westernised populations spanning multiple countries and continents compared to Westernised populations is remarkable, with the only exception being from Mongolia (Liu et al., 2016). The Mongolian cohort sampled both urban dwellers and rural non-Westernised populations. While the urban dwellers have a prevalence closer to that of the non-Westernised populations (Clade A prevalence; 100%, B; 73.3%, C; 80%, D; 42.2%), this is still generally lower than the rural non-Westernised Mongolian population (Clade A prevalence; 98.5%, B; 76.9%, C; 84.6%, D; 56.9%). Although *P. copri* was observed at a much lower abundance in Westernised populations, all four clades were detected, and among those Clade A was the most prevalent type (Figure 3A).

**Figure 3.**
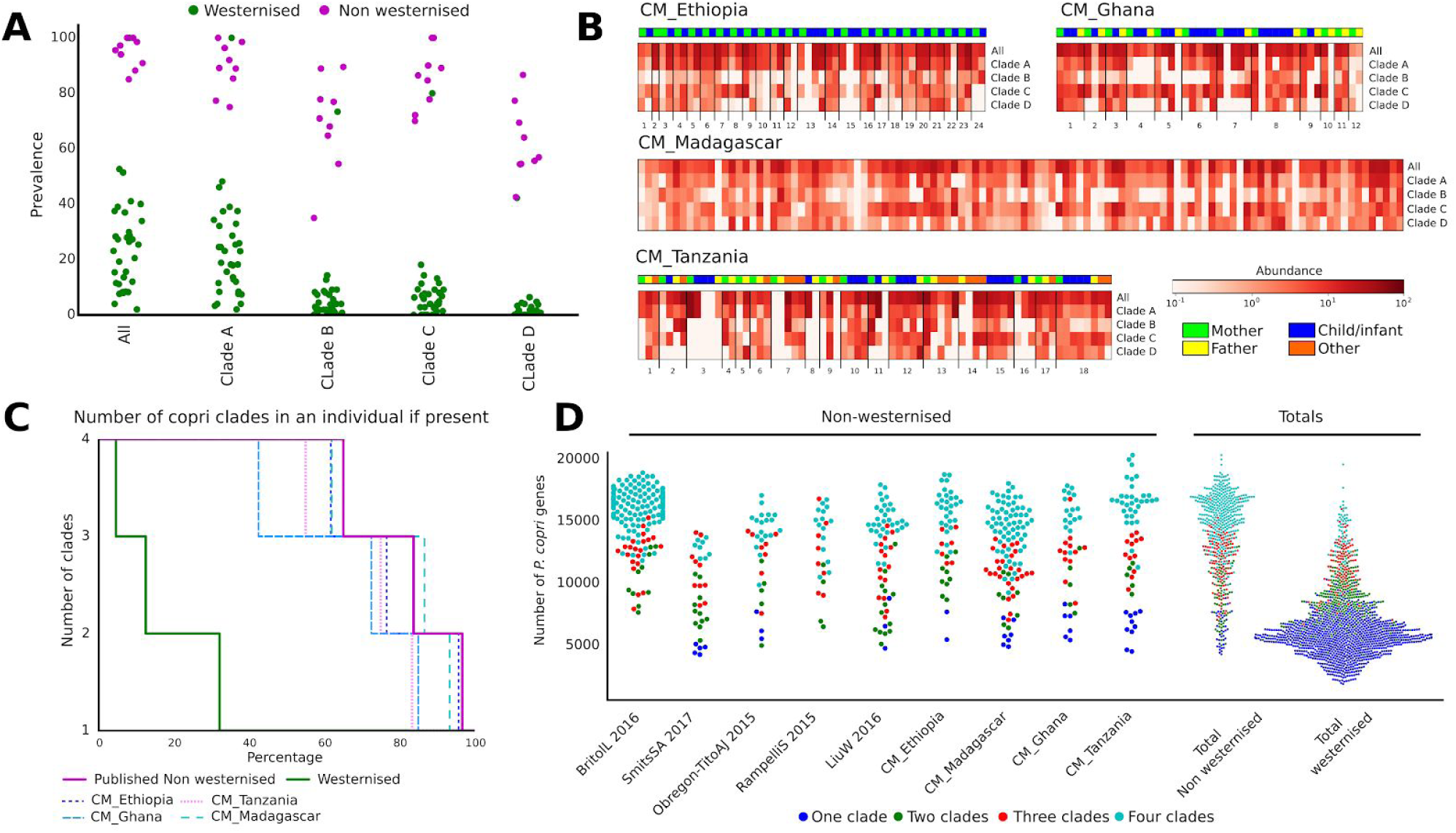
Prevalence of the *P. copri* complex and its association with non-Westernised populations. **A)** *P. copri* prevalence in individual non-Westernised and Westernised datasets, “All” refers to the prevalence of any of the four clades being present. **B)** Prevalence and abundance of the *P. copri* complex in the four newly sequenced non-Westernised datasets. **C)** Percentage of individuals harbouring multiple *P. copri* clades. **D)** *P. copri* pangenome sizes for non-Westernised individuals by dataset compared to Westernised individuals.

Considering the high prevalence of the four *P. copri* clades in non-Westernised populations we next sought to identify if these clades are mutually exclusive or able to co-inhabit in the intestine. Analysis of our newly sequenced datasets clearly displayed multiple clades within individuals (Figure 3B), which confirms the results observed when analysing all the other available datasets (Figure 3C). Strikingly, for the 95.4% of non-Westernised individuals with at least one *P. copri* complex clade, in 61.6% of these all four clades were detectable, in 82.0% at least three, and at least two in 93.8% of individuals. The high-percentage of individuals carrying multiple clades was a consistent feature observed across all non-Westernised datasets, spanning four continents (Figure 3C). In comparison, for the smaller fraction of 29.6% of Westernised individuals with at least one *P. copri* complex clade, only 4.6% had all four clades, 12.5% at least three and 32.1% more than one. Therefore, we demonstrate that not only is *P. copri* prevalence higher in non-Westernised populations but the pattern of multi-clade co-presence in these populations is also a defining characteristic.

Because of the existence of multiple clades within an individual (Figure 3B **and** 3C), and the observation of sizeable inter-clade diversity based on gene content (Figure 1D), we decided to estimate the sum of unique *P. copri* complex genes within each individual, or rather “the within individual *P. copri* pangenome” (see **Methods**). As expected, individuals with multiple clades tended towards a larger number of unique *P. copri* genes (Figure 3D), and as multiple clades is a feature of being non-Westernised, a considerably larger *P. copri* functional potential was revealed in these populations (Figure 3D, **Supplementary Figure 7**).

### Evidence of distinct carbohydrate metabolism repertoires in the four *P. copri* complex clades

To investigate the potential functional diversity of the *P. copri* complex, we first annotated the open reading frames (ORFs) for each genome using the eggNOG database (Huerta-Cepas et al., 2017) (see **Methods**). Between and even within the four clades of the complex we observed considerable diversity, with Clade B being the most dissimilar based on the overall distance of the eggNOG functional profiles (Figure 4A), which is consistent with the inter-clade genetic diversity discussed above (Figure 1B-D). Some of the distinguishing functionalities included sulphur metabolism and assimilation, which were enriched in all clades relative to B (**Supplementary Table S5**). Similarly, in carbohydrate metabolism, beta-galactosidase was found to be absent in Clade B while being prevalent in all other clades (at least >60% prevalence). In the metabolism of cofactors and vitamins, genes responsible for folate metabolism were depleted in Clade D. Interestingly, Clade D also had the least diversity of antimicrobial resistance genes lacking 5 out of 7 identified as differentially prevalent in the other three clades. Differences were also noticeable in membrane transporters; for instance, the polyamine spermidine/putrescine ABC transporter (pot*ABCD*) was present in almost all members of Clade A, C and D but never observed in Clade B. Conversely, an energy coupling factor (ECF) type ABC transporter that could be responsible for micronutrient uptake was solely found in a subset of genomes of Clade B (27% of genomes).

**Figure 4.**
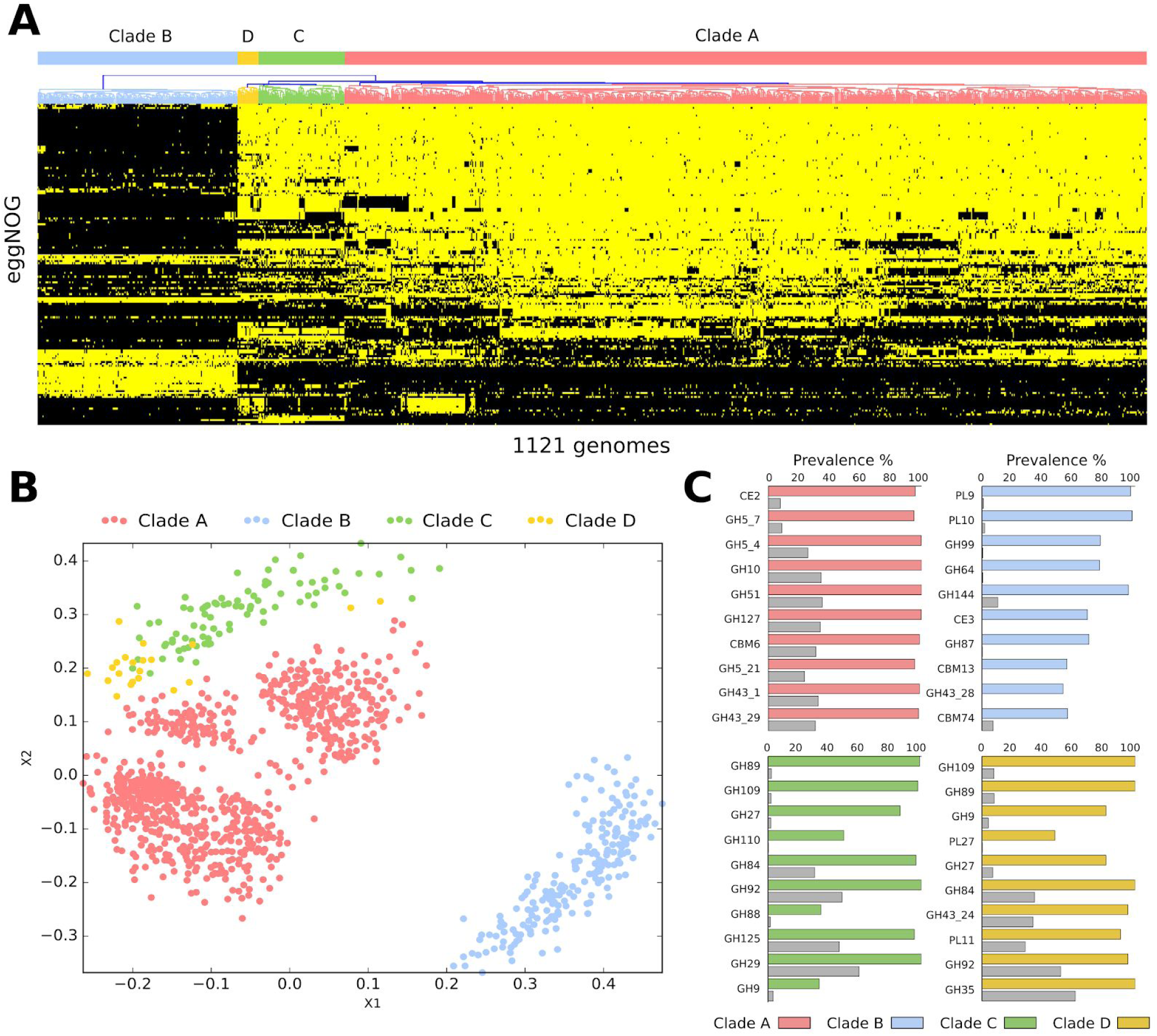
Functional diversity of the *P. copri* complex. **A)** Presence and absence of eggNOG functions significantly different between the four *P. copri* clades (yellow present, black absent) (see **Methods**). **B)** Multidimensional scaling (MDS) ordination based on CAZy families present in each genome showing distinct inter and intra clustering in the *P. copri* complex. **C)** The ten most enriched CAZy families for each clade compared to all other clades (grey bars). For full list of CAZy prevalence see Supplementary **Table S6**.

One reported feature of *P. copri* is its effect on glucose homeostasis (De Vadder et al., 2016; Kovatcheva-Datchary et al., 2015; Pedersen et al., 2016), with one recent study suggesting a positive benefit via succinate production (De Vadder et al., 2016). While potential succinate production was observed in all clades the genes responsible were less prevalent in Clade B (p-value < 5.2e-37, Bonferroni-corrected Fisher-exact test). On the contrary, high-levels of circulating branched chain amino acids (BCAA), linked to the development of insulin resistance, have been associated to higher levels of *P. copri* in the gut microbiome (Pedersen et al. 2016). In addition, the prevalence of genes for BCAA biosynthesis in the *P. copri* pangenome has been recently shown to be diet-dependent and associated to actual urinary BCAA levels (De Filippis et al., 2019). Here we found that BCAA biosynthesis genes were widespread across all four clades of the *P. copri* complex, with >85% prevalence in all cases.

The *P. copri* complex is strongly associated with non-Westernised populations which have diets that tend to be higher in fibre and complex carbohydrates and lower in fats and animal protein compared to typical Western diets (De Filippo et al., 2010; Segata, 2015; Statovci et al., 2017). To specifically look at the *P. copri* complex for potential carbohydrate utilisation, the genomes were also screened for carbohydrate active enzymes (CAZymes) (Lombard et al., 2014) (see **Methods**). While many of these CAZy families were found to be prevalent across all four *P. copri* clades (**Supplementary Table S6**), considerable variability in the presence of these families was seen between and even within the different clades (Figure 4B, 4C and **Supplementary Figure S8**). To focus on families potentially associated with plant-derived carbohydrate degradation (e.g. cellulose, hemicellulose and pectin) each family was ascribed a broad substrate specificity via manual curation (**Supplement Figure S9**).

While all clades were found to have the potential to degrade plant-derived carbohydrates, not all CAZy families were represented or equally distributed throughout the four clades (**Supplementary Figure S9);** for example, the polysaccharide pectin-degrading families PL9 and PL10 were highly prevalent and nearly exclusively present in Clade B. In Clade D, the GH9 CAZy family of cellulases was particularly enriched compared to the other clades (Figure 4C). The distinct clustering based on CAZy gene content (Figure 4B) displaying inter- and intra-clade functional differences suggests overlapping but potential heterogeneity in carbohydrate metabolism. The frequent co-presence of all four clades in non-Westernised populations would suggest they are non-competing and therefore niche separated. While it cannot be discounted that these four clades are spatially separated in the intestine, the ability to utilise a differing array of carbohydrates or substrate specificities could potentially be the driver of this separation. Within an individual the presence of multiple clades collectively offers a larger and perhaps complementary functionality to efficiently metabolise a wide range of dietary fibre.

### *P. copri* diversity in ancient human gut contents resembles that of non-Westernised populations and gives insights into its evolutionary history

To test whether the high *P. copri* prevalence and co-presence of the four clades of the complex in non-Westernised populations reflects the composition in ancient human gut microbiomes, we analysed four archaeological gut contents for the presence of *P. copri*. We studied material from the lower intestinal tract and lung tissue of the Iceman, a 5,300-year-old natural ice mummy (Spindler, 1994). The Iceman genetically belongs to the Early European Farmers and originated and lived in Southern Europe, in the Eastern Italian Alps (Haak et al., 2015; Keller et al., 2012; Lazaridis et al., 2014; Müller et al., 2003). (Figure 5A). We also analysed three coprolite samples (fossilized faeces) (Figure 5A) recovered from the pre-Columbian (1300 ± 100 B.P.) site “La Cueva de los Muertos Chiquitos” from Durango, a Northwestern state of Mexico (Brooks et al., 1962) (see **Methods**).

**Figure 5.**
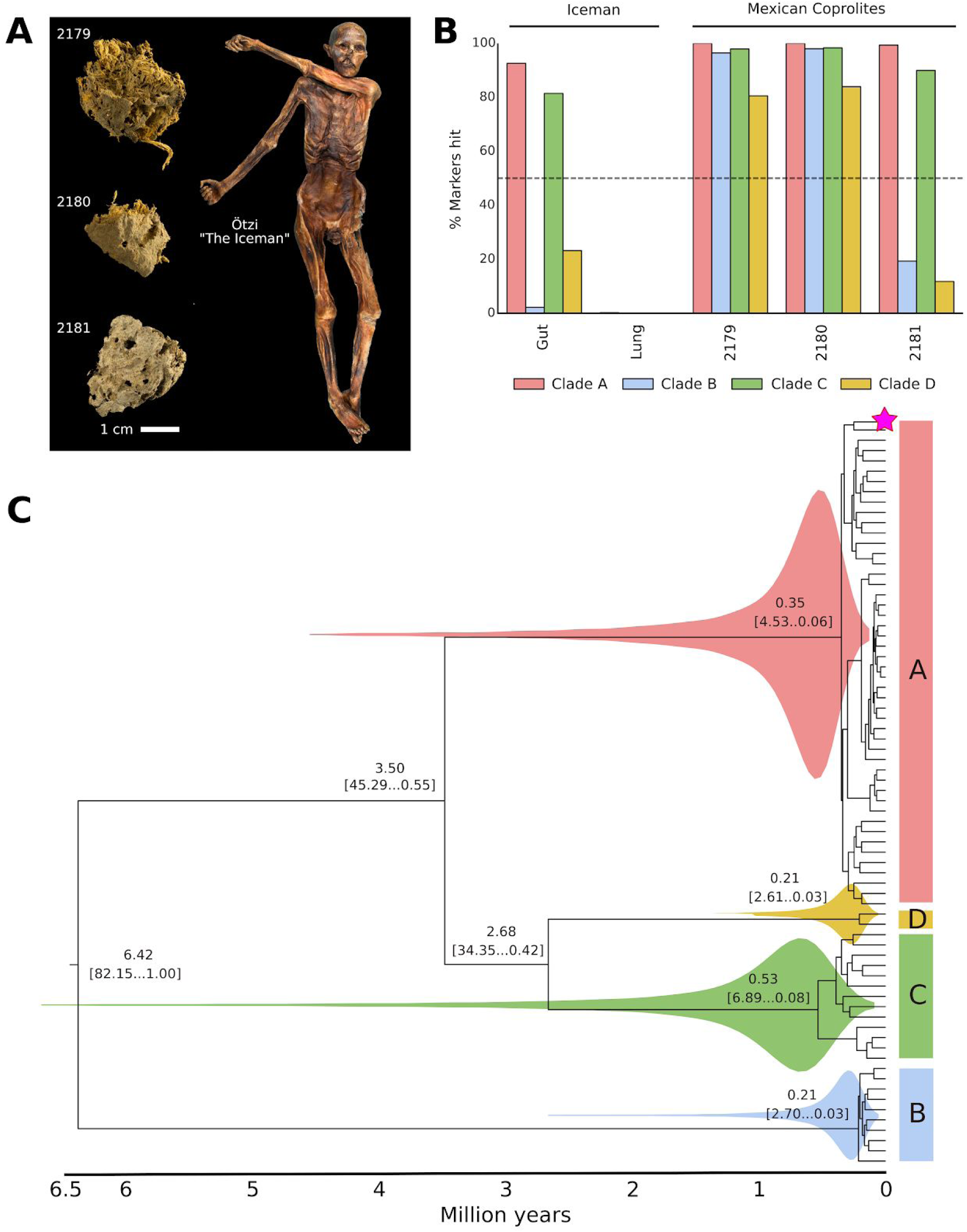
Ancient microbiomes and the evolutionary history of the *P. copri* complex. **A)** Ancient Mexican coprolite samples and intestinal and lung tissue sampled from the Iceman, a natural ice mummy. **B)** Percentage of positive *P. copri* clade specific markers identified in each ancient metagenomic sample. **C)** Time measured phylogenetic tree of the *P. copri* complex, magenta star indicates ancient coprolite sample, 2180 (see **Methods**).

We found the *P. copri* complex to be prevalent both in the Mexican and the European ancient gut metagenomes (Figure 5B). All samples had at least two *P. copri* clades (Clades A and C) and in two coprolites all four clades could be detected. The higher prevalence of clade A and C in our ancient samples mirrors the tendency of modern day populations (both Westernised and non-Westernised) where these two clades are more prevalent (Figure 3A). To discount the possibility of a non-ancient gut origin, we verified that the *P. copri* reads displayed damage patterns indicative of ancient DNA (**Supplementary Figure 10**) (Orlando et al., 2015) and in a control sample (Iceman lung tissue) no *P. copri* clades were detected (positive for only a single marker of the 2,448 *P. copri* complex specific markers) (Figure 5B). Two characteristics point toward a similarity between the ancient samples and modern non-Westernised populations. First, *P. copri* is highly prevalent in the ancient samples as in non-Westernised samples. Second, the ancient samples are characterised by a high clade co-presence (presence of 2 to 4 distinct clades) as observed in non-Westernised individuals. Although we have analysed only a small number of ancient metagenomes, these observations suggest that the *P. copri* carriage in non-Western populations is more akin to our ancestors.

To calibrate a *P. copri* phylogeny we screened all ancient samples and found that one coprolite (sample 2180, radiocarbon dated A.D. 673 to 768, see **Methods**) had sufficiently high coverage of Clade A for clocking (see **Methods**). Model selection indicated that this dataset is best modelled by a strict clock suggesting a constant rate of evolution through time and in different *P. copri* clades (see **Methods**). All our divergence estimates converged satisfactorily on clear posterior means (Figure 5C) and the age estimates indicate that *P. copri* began to diversify (split of clade B) ~6.5 million years ago (Figure 5C). This split is concurrent with the proposed divergence of human and chimpanzees (Patterson et al., 2006a). The diversification of clades A, D and C is estimated to have occured between ~3.5 and ~2.5 million years ago when the first archaic species of the genus *Homo* developed (Ungar and Sponheimer, 2011; Wood and Collard, 1999). The differentiation within each of the clades is instead relatively recent with the median estimates following that of the emergence of *Homo sapiens* circa 315 ka (Hublin et al., 2017). Despite the range in estimated clade divergence, even at the lowest estimation (420 ka) this occurred well before the first human migration waves out of Africa circa 90 to 194 ka years ago (Grün et al., 2005; Hershkovitz et al., 2018). This would indicate that the four clades of the *P. copri* complex were a feature of our pre-migratory human ancestors.

Further support of *P. copri* clade diversification prior to migration is that the high prevalence of all four clades and multi-clades within an individual is a consistent feature of disparate non-Westernised populations in Africa, Oceania, South America and Asia. This observation in non-Westernised populations that are geographically and socially restricted would suggest vertical rather than horizontal acquisition of *P. copri* between the populations. This, together with the estimation of clade divergence, implies the *P. copri* complex has been a long-standing feature of the human microbiome. This analysis suggests the underrepresentation of *P. copri* in Westernised populations is the consequence of its loss in response to Westernisation, which has been rapidly occurring in an almost infinitesimal time frame relative to host-microbe coevolution.

## Discussion

We demonstrate that *P. copri* is not a monotypic species but four clearly defined clades, each spanning a diversity that is typical of distinct species (Figure 1), and all four clades have the potential to reside either solely or in combination within an individual (Figure 3C). We propose to name this group the *P. copri* complex, comprising clades A, B, C and D. The insights that we have gained into the *P. copri* complex genetics and population genomics relied on isolate sequencing, on new sequencing from underrepresented non-Westernised populations, and largely on the tremendous resource of publicly available metagenomic datasets, covering multiple geographies, diseases and lifestyles. This led to the observation that the *P. copri* complex is globally distributed (Figure 2), but with a highly structured distribution, both in terms of prevalence and the presence of multiple clades within an individual, with non-Westernised populations (Figure 3).

While we concede the term Westernised versus non-Westernised serves to demarcate what may be better seen as a continuum along multiple lifestyle parameters, the distinction nonetheless has its merits, as interest grows in comparing our Westernised microbiomes to those presumed more akin to our ancestral microbiomes. Recent studies have expanded our understanding of the microbial diversity of non-Westernised populations (Hansen et al., 2019; Pasolli et al., 2019), and the rapid loss of diversity with Westernisation (Vangay et al., 2018). What is still to be determined is the consequences of this microbial impoverishment with respect to the wider gut microbial ecosystem and its impact on human health.

Evidence from the analysis of ancient stool samples (Figure 5) points towards *P. copri* diverging into four clades prior to the first human migration events out of Africa. The fact that we consistently observe high prevalence in globally disparate non-Westernised populations and in ancient microbiomes suggest the loss of *P. copri* might be a result of Westernisation. A major element of Westernisation has been a shift in diet over the course of the last two centuries with the advent of industrialisation and food processing, from one high in fibre and complex carbohydrates to one high in sodium, fat, and simple sugars and low in fibre. It was previously shown that *P. copri* provides a host benefit in response to a high-fibre diet (De Vadder et al., 2016; Kovatcheva-Datchary et al., 2015), but not one high in fat (Pedersen et al., 2016). The *P. copri* complex shows a diversity in plant-derived carbohydrate utilisation (Figure 4) which may also suggest diet is a key driver responsible for its ultimate demise in Westernised populations.

Diet in particular seems to play a pivotal role in the case of *P. copri*, yet it is extremely difficult to study this influence in the context of long-term human dietary modifications spanning multiple generations. Given previous work associating multi-generational microbial impoverishment with dietary changes in mice (Sonnenburg et al., 2016), clearly more work is required both *in silico* and using *in vitro* studies to functionally associate and characterise the *P. copri* complex with respect to long-term dietary exposures, transmission, and retention. In part, *P. copri* has come to attention based on its association with disease, but in this study we found no clear evidence that particular clades are associated with the subset of health conditions available for meta-analysis. Nevertheless, it cannot be disregarded that such associations may exist, possibly only at the sub-clade level, but such an investigation would require the power of a far larger number of disease-specific cohorts than are currently available.

Finally, it is particularly notable that the analysis approach taken here is generalisable to other microbial species in instances where there are minimal reference isolate sequences available. This is in principle the case for species that are understudied due to being recalcitrant to cultivation or because they have not been the focus of sequencing efforts. The ever-increasing number of publicly available metagenomes will serve this end, as well as likely add clarity to whether *P. copri* is considered either a positive or negative influence on health in the context of other microbiome members, diet, lifestyle and host genetic factors. This study reveals *P. copri* is far more complex than previously imagined and it will be important in future studies to appreciate this in order not to oversimplify and underestimate the potential *P. copri* diversity within the human gut microbiome.

## Material and Methods

The overall aim of the study was to identify and recover *P. copri* genomes in publicly available intestinal metagenomic datasets and new non-Westernised datasets presented in this work. These datasets represent multiple countries, host-conditions and lifestyles. The process involved collating a panel of 72 high-quality *P. copri* genomes using manually guided metagenomic genome binning and sequencing new *P. copri* isolates and publicly available reference sequences. The resulting panel of genomes was used to automatically recover additional *P. copri* genomes from single-sample assembled metagenomes. In addition, clade-specific markers were designed to accurately identify the prevalence and abundance of the *P. copri* clades across datasets.

### Metagenomic assembly

Metagenomic samples were assembled using metaSpades (Version 3.10.1) (Nurk et al., 2017) using default parameters, chosen due to its reported performance compared to other assemblers (Forouzan et al., 2018; Pasolli et al., 2019; van der Walt et al., 2017). Samples that exceeded permitted memory requirements (>1Tb of RAM) or those for which only unpaired reads were available were assembled using Megahit (Version 1.1.1) (Li et al., 2015) using default parameters. Only assembled contigs ≥ 1kb were considered further.

### Constructing a *P. copri* genome panel with newly sequenced isolates and manually curated genomes from metagenomes

A panel of 72 *P. copri* genomes were collated consisting of two publicly available reference genomes (RefSeq assembly accessions: GCF_000157935.1, GCF_002224675.1), 15 newly described isolate genomes (see below) and 55 manually curated metagenomically assembled genomes.

*P. copri* strains were isolated from stool from healthy subjects and new onset rheumatoid arthritis patients. Stool was collected into anaerobic transport media (Anaerobe Systems), then streaked on BRU and LKV plates (Anaerobe Systems). After 24-48h, individual colonies were picked and screened with *Prevotella*-specific PCR primers, and the 16S rRNA V3-V4 sequence was confirmed by Sanger sequencing (Fehlner-Peach et al., manuscript in preparation). *Prevotella*-positive isolates were grown on BRU plates, and mature colonies were collected for genomic DNA isolation with the DNeasy PowerSoil Kit (Qiagen). Libraries were prepared for sequencing on the HiSeq2500 platform with the TruSeq DNA PCR-free Library Prep Kit (Illumina). In total 83 *P. copri* isolates were sequenced. As this included multi-sampling from the same individual, 15 isolates were selected for this study that represented the total genetic diversity of the isolate dataset.

The 55 genome bins were recovered using anvi’o (Version 2.3.2) (Eren et al., 2015) applied on a set of assembled metagenomes. Anvi’o provides a platform for metagenomic genome binning and offers the ability to manually asses and curate those bins, potentially increasing accuracy compared to automated binning methods, but at the expense of being low throughput. Briefly, the 100 metagenomic samples determined to have a high abundance of *P. copri* based on MetaPhlan2 (Truong et al., 2015) were selected. The metagenomic samples were assembled (see above) and reads mapped back to the contigs using Bowtie2 (version 2.2.9, using “very-sensitive-local” parameter) (Langmead and Salzberg, 2012). Contigs (>2.5kb) were clustered by anvi’o based on coverage and tetranucleotide frequency, and manually curated. All recovered bins were subjected to strict quality control (see below), resulting in 55 high-quality genome bins.

### Automated recovery of *P. copri* genomes from >6500 metagenomes

To automatically recover *P.copri* genomes from metagenomes, we first assembled each metagenome (see above) and for each assembly its contigs were mapped against the panel of 72 high-quality reference genomes representing the four clades of the *P. copri* complex (described above) using Blast (version 2.6.0+) (Altschul et al., 1990). Only contigs with a nucleotide identity ≥95% and an alignment ≥50% were considered further and placed into one of the four *P. copri* bins (Clade A, B, C or D) based on the membership of the reference genome. On the rare occasion a contig was ≥95% identical and aligned over ≥50% to multiple reference genomes representing different clades, the contig was placed into a single clade bin based on best BitScore, if this score was ≥10% than any other competing clade(s). If the BitScore threshold was not satisfied, the contig could not confidently be placed and was not considered further. All recovered *P. copri* metagenomic genome were assessed for quality (see below).

### *P. copri* genome quality control

All *P. copri* genomes were strictly quality controlled. QC involved four steps 1) genome size 2) estimated completeness, 3) estimated contamination and 4) level of strain heterogeneity. Only genome bins >2.5 Mb <5.0 Mb and composed of <500 contigs were considered. CheckM (Parks et al., 2015) was used to estimate the completeness and level of contamination. High quality genomes were those >95% completeness, <5% contamination, except for *P. copri* clade D where a completeness of >90% was used to be more inclusive. For the newly sequenced non-Westernised datasets (see section: westernisation and additional Non-Westernised datasets, below) and the manually curated metagenomically assembled genomes a threshold of >90% completeness was also selected. We also investigated strain-level diversity for each of the *P. copri* clades within a sample as this could indicate contig chimeric assembly. Strain-level heterogeneity was estimated using an in-house developed tool, CMseq, available here: https://bitbucket.org/CibioCM/cmseq/commits/41082ef. Firstly, protein coding genes of the contigs were predicted with prodigal (Hyatt et al., 2010) implemented in the prokka pipeline (Version 1.11) (Seemann, 2014). To avoid overestimating strain-heterogeneity due to genes in common across the four *P. copri* clades, only the clade specific genes (see below) were considered as a proxy to estimate strain heterogeneity. Secondly, metagenomic reads were mapped to the assembled contigs using Bowtie2 (Langmead and Salzberg, 2012) (version 2.2.9, using “very-sensitive-local” parameter) and for each coding nucleotide base calls were only considered if there was >10X coverage and a PHRED quality score of ≥ 30. Each position was considered non-polymorphic if the frequency of the dominant allele was >80%. When calculating the overall contig polymorphic rate only the non-synonymous positions were considered.

### Genetic distance between and within the *P. copri* complex and related species

Average nucleotide distance (ANI) pairwise were calculated using pyani (version 0.2.6; option ‘-m ANIb’) (Pritchard et al., 2015) considering representative subset of *P. copri* genomes of the four clades (25 for clade A, B and C, and all the 15 genomes of clade D) and publicly available reference genomes of the *Prevotella*, *Alloprevotella* and *Paraprevotella* genera available from NCBI RefSeq. Distances scores were filtered to include only pairwise comparison where alignment lengths exceeded 500,000 bp. Pairwise core genome distances based on comparisons of single nucleotide polymorphisms were calculated on the core genome alignment of all 1,023 *P. copri* genomes. The core genome alignments were produced utilising PRANK (Löytynoja, 2014) as part of the Roary pipeline (version 3.11) (Page et al., 2015) with the parameters of 90% similarity identity for gene clustering and present in 90% of genomes for defining core genes. The pangenome-based matrix also produced from Roary was used to compare the pairwise gene content similarity calculated using the Jaccard similarity coefficient as part of the SciPy package (Bressert, 2012). To infer instances of strain sharing between individuals, normalised phylogenetic distances on the *P. copri* phylogeny were compared and called as the same strain based on a 0.2% identity threshold as previously described (Truong et al., 2017).

### Phylogenetic analysis

The phylogenetic analyses were performed with PhyloPhlAn (Segata et al., 2013) using the new version available in the “dev” branch of the repository (commit 7c38e19, https://bitbucket.org/nsegata/phylophlan).

The phylogeny in Figure 1A was built using the 400 universal marker genes as identified by PhyloPhlAn using the following parameters: “--diversity low --fast”. The set of external tools with their respective options is reported below:

- Diamond version v0.9.9.110, (Buchfink et al., 2015), with “blastx” for the nucleotide-based mapping, “blastp” for the amino-acid based mapping, and “--more-sensitive --id 50 --max-hsps 35 -k 0” in both cases
- MAFFT version v7.310, (Katoh and Standley, 2013), with “--localpair --maxiterate 1000 --anysymbol --auto” options
- trimAl version 1.2rev59, (Capella-Gutiérrez et al., 2009), with “-gappyout” option
- FastTree version 2.1.10, (Price et al., 2010), with “-mlacc 2 -slownni -spr 4 -fastest -mlnni 4 -no2nd -gtr -nt” options
- RAxML version 8.1.15 (Stamatakis, 2014), with “-p 1989 -m GTRCAT -t <FastTree phylogeny>” options

The tree was built on a total of 90 genomes comprising a subset of 25 representative genomes for the three clades A, B and C whereas for clade D all 15 genomes were considered.

The phylogeny in **Supplementary Data File S1** is based on the 210 set of core genes screened to be monophyletic as described in the section below on molecular dating. The phylogeny has been reconstructed using PhyloPhlAn with the following parameters: “--diversity low --trim greedy --remove_fragmentary_entries”. Additionally, the set of external tools with their options is reported below:

- Blastn version 2.6.0+, (Altschul et al., 1990), with “-outfmt 6 -max_target_seqs 1000000” options
- MAFFT, trimAl, FastTree, and RAxML were run with the same options as reported above.

The phylogenies in Figure 2 and in **Supplementary Figure S6** were built with PhyloPhlAn using the set of core genes for each *P. copri* clade (>95% prevalent across all genomes within a clade) determined using Roary (version 3.11) (Page et al., 2015) with a minimum gene identity of 90%. PhyloPhlAn was run using the following parameters: “--mutation_rates --min_num_entries <97% of the number of input genomes> --diversity low”. The set of external tools used is: Blastn, MAFFT, trimAl, FastTree, and RAxML, and they were executed with the same options as reported above. The phylogenetic trees in Figure 2, and **Supplementary Figure S6** were visualized using GraPhlAn (version 1.1.3, (Asnicar et al., 2015)).

### The *P. copri* pangenome and evaluating prevalence and abundance

The protein coding regions for the 72 *P. copri* genome panel (see above) were predicted using Prodigal (Hyatt et al., 2010) as part of the prokka pipeline (Version 1.11) (Seemann, 2014) and the total *P. copri* pangenome determined using Roary with 90% similarity identity parameter (Version 3.11) (Page et al., 2015). Markers specific to each clade of the *P. copri* complex where defined as present in >95% of the *P. copri* genomes of a given clade but absent in all others. This gave for Clade A n=430 markers, for Clade B n=954, for Clade C n=479 and for Clade D n=585. To determine if a *P. copri* clade is present in a metagenomic sample reads were mapped to the clade specific markers using Bowtie2 (Langmead and Salzberg, 2012) and mappings processed using PanPhlAn (Scholz et al., 2016). A marker was scored present if had a coverage ≥0.5X, and a clade present if ≥50% of the clade specific markers were hit. Estimation of *P. copri* clade relative abundance was calculated thus: (Mean clade marker coverage × approximated genome size(bp)) / total metagenome size (bp).

### Westernisation and additional Non-Westernised datasets

Westernisation as the adoption of a Westernised lifestyle and culture can trace its origins to industrialisation and its promotion of urbanisation over the past two centuries. Westernisation has had a profound effect on human populations, due to access to healthcare and pharmaceutical products, hygiene and sanitation, changes in diets (processed, high-fat, low in complex carbohydrates but rich in refined sugars and salt), population density increase and reduced exposure to livestock. Westernisation is nonetheless difficult to ascribe as it demarcates populations which are clearly on a continuum. In this study definition of “Westernised” and “non-Westernised” is considered based on how populations differ based on the above criteria and how the samples were reported in the original publication.

In this study five previous studies were considered where non-Westernised populations have been sampled from Fiji (Brito et al., 2016), Peru (Obregon-Tito et al., 2015), Tanzania (Rampelli et al., 2015; Smits et al., 2017) and Mongolia (Liu et al., 2016). In addition, we recently sequenced a population of adults from a rainforest region in North-eastern Madagascar (110 metagenomes) (Pasolli et al., 2019). We expanded upon a dataset of 5 samples sequenced from an established cohort in Ethiopia from Gimbichu in the Oromia region (Pasolli et al., 2019) with 45 additional samples. This cohort included 24 mothers and their infant(s) for a total of 50 metagenomes. We also sequenced two new non-Westernised populations from Ghana and Tanzania. In Ghana 12 extended families from the Asante Akim North district region were sampled where the local occupation is subsistence farming and the wider economy based on farming cash crops such as cocoa and plantain and there is also a commercial poultry industry. (44 metagenomes). From Tanzania samples were collected from 18 families from Korogwe District region where local employment and economy is based on agriculture particularly based on sisal fibres, cashew nuts and cotton (68 metagenomes). For all samples DNA was extracted using the PowerSoil DNA isolation kit (MoBio) as previously described (Human Microbiome Project Consortium, 2012). Libraries were constructed using the NexteraXT DNA Library Preparation Kit (Illumina) and sequenced on the Illumina HiSeq2500 100nt paired end platform with a target depth of 5Gb/sample.

### Genome functional potential analysis

We performed the functional annotation using the EggNOG mapper (version 1.0.3) (Huerta-Cepas et al., 2017) that is based on the EggNOG orthology system (Huerta-Cepas et al., 2016) and the sequence searches performed using HMMER (Eddy, 2011). We used the KEGG Brite Hierarchy to screen the EggNOG annotations that are shown in Figure 4A. In this figure, for each clade, we report only the eggNOGs that are significantly different in each of its three pairwise comparisons to the other clades (p-value < 0.01, Bonferroni corrected Fisher-exact test). CAZy enzymes (Lombard et al., 2014) (http://www.cazy.org/) were predicted with HMMSEARCH (Version 3.1b2) (Eddy, 2011) against dbCAN HMMs v6 using default parameters and applying post-processing stringency cut-offs as suggested (Yin et al., 2012).

### Iceman samples and Mexican coprolite material

In this study we metageomically analyzed archaeological gut contents for the presence of *P. copri*. The analyzed material includes gut content and lung tissue (negative control) of the Iceman, a European Copper Age ice mummy (Figure 5). The Iceman, commonly referred to as “Ötzi”, is one of the oldest human mummies discovered. His body was preserved for more than 5,300 years in an Italian Alpine glacier before he was discovered by two German mountaineers at an altitude of 3,210 m above sea level in September 1991. The mummy is now conserved at the Archaeological Museum in Bolzano, Italy, together with an array of accompanying artifacts (www.iceman.it). The Iceman was naturally mummified by freeze-drying (Lynnerup, 2007). Therefore, his body tissues and intestines still contain well preserved ancient biomolecules (DNA, proteins, lipids) that allowed e.g. the reconstruction of the Iceman’s genome (Keller et al., 2012), the genomic analysis of the stomach pathogen *Helicobacter pylori* (Maixner et al., 2016), and the molecular reconstruction of the Iceman’s last meal (Maixner et al., 2018). In addition, we subjected three ancient coprolite samples from a Mexican cave to metagenomics analysis (Figure 5). The archaeological site “La Cueva de los Muertos Chiquitos” in the northern Durango region of el Zape, Mexico, was excavated by Brooks and colleagues in the early 1960s (Brooks et al., 1962). The sub-humid climate in this natural cave at an altitude of approx. 1,800 m above sea level provided favourable conditions for the preservation of various ancient remains including human skeletons, botanical artefacts, quids and coprolites. The site was dated by previous radiocarbon dating of a single wood sample from one of the oldest levels (square B4, level 24-28) to A.D. 600 (1300 ± 100 B.P.) (Brooks et al., 1962). The ancient remains have been previously subjected to botanical (Brooks et al., 1962), dietary (Hammerl et al., 2015; Meade, 1994), parasitological (Camacho et al., 2018; Jiménez et al., 2012; Morrow and Reinhard, 2016), and molecular analysis (Tito et al., 2008, 2012). All three coprolite samples used in this study were discovered in square B4 in two different levels (**Supplementary Table S7**). The samples were stored in the Pathoecology Laboratory in the School of Natural Resources at the University of Nebraska-Lincoln in Lincoln, Nebraska. We obtained radiocarbon dates at the Curt-Engelholm-Centre for Archaeometry, Mannheim, Germany for coprolite sample 2180 from level 16-20 (A.D. 673 to 768, 1,284 ± 16 B.P.) that confirms the previous direct dating of the pre-Columbian archaeological site (**Supplementary Table S8**).

The molecular analysis of the Iceman samples and of the ancient human coprolites was conducted at the ancient DNA laboratory of the EURAC Institute for Mummy Studies in Bolzano, Italy. Sample preparation and DNA extraction was performed in a dedicated pre-PCR area following the strict procedures required for studies of ancient DNA: use of protective clothing, UV-light exposure of the equipment and bleach sterilization of surfaces, use of PCR workstations and filtered pipette tips. DNA was extracted from the archaeological specimen using a chloroform-based DNA extraction method according to the protocol of (Tang et al., 2008). Libraries for the sequencing runs were generated with a modified protocol for Illumina multiplex sequencing (Kircher et al., 2012; Meyer and Kircher, 2010). Libraries of the Mexican coprolite samples were sequenced on Illumina HiSeq2500 platforms using 101–base pair paired-end sequencing kits. The Iceman samples were sequenced on an Illumina HiSeqX platform using the 150–base pair paired-end sequencing kit.

Paired Illumina reads were quality-checked and processed (adapter removal and read merging) as previously described in (Maixner et al., 2018). Reads were mapped using Bowtie2 (Langmead and Salzberg, 2012) to the human genome (build Hg19, default mapping parameters) (Rosenbloom et al., 2015), the human mtDNA reference genome (rCRS, mapping parameter --very-sensitive-local) (Andrews et al., 1999), and selected *P. copri* genomes from the four clades. For details to the mapping results please refer to **Supplementary Table S9**. To deduplicate the mapped reads we used the DeDup tool (https://github.com/apeltzer/DeDup). The minimum mapping and base quality were both 30. The resulting bam files were used to check for characteristic aDNA nucleotide misincorporation frequency patterns using mapDamage2 (Jónsson et al., 2013). Both human and bacterial reads display low but already increased frequencies of C to T substitutions close to the fragment ends characteristic of ancient DNA (Orlando et al., 2015) (**Supplementary Figure S10**). Reads of the Iceman lung tissue metagenome that mapped to one single marker of the 2,448 *P. copri* complex specific markers display no DNA damage. The sex of the mapped human reads was assigned using a Maximum likelihood method, based on the karyotype frequency of X and Y chromosomal reads (Skoglund et al., 2013) (**Supplementary Table S9**). Estimation of human contamination rates using Schmutzi (Renaud et al., 2015) and ANGSD (Korneliussen et al., 2014) was in most samples not possible due to low damage pattern rates in the mitochondrial reads and due to the low coverage of the X chromosome (sample 2180), respectively. The Iceman samples with sufficient X-chromosome coverage show low contamination in the autosomal DNA when using ANGSD (**Supplementary Table S9**). Analysis of the human mitochondrial and autosomal variants provided further evidence for the sample origin and authenticity of the data. Variants in the mitochondrial genome were called using SAMtools mpileup and bcftoools (Li et al., 2009) with stringent filtering options (quality>30). Visual inspection of the called variants identified only less than 1% low-frequency variants that could be indicative for contamination.The haplogroup was identified by submitting the variant calling file to the HaploGrep website (Weissensteiner et al., 2016) (**Supplementary Table S9**). The human mitochondrial genomes in both Iceman samples carry the same variants as reported in previous Iceman genomic studies and belong to the K1f haplotype (Ermini et al., 2008; Keller et al., 2012). Importantly, in the Mexican coprolite samples 2179 and 2180 both detected mitochondrial haplogroups (C1b and B2) belong to the four main pan-American mtDNA lineages (Achilli et al., 2008; Bodner et al., 2012; Perego et al., 2010; Tamm et al., 2007). Furthermore, both haplogroups have been detected in previous studies in ancient human remains from Meso- and South America (Fehren-Schmitz et al., 2015; Gómez-Carballa et al., 2015; Llamas et al., 2016; Posth et al., 2018; Tackney et al., 2015) and haplogroup C1b has still nowadays its highest frequency in Peru and Mexico (Gómez-Carballa et al., 2015). We extended our analysis to the human autosomal data and called pseudodiploid genotypes using SAMtools mpileup (Li et al., 2009) and PileupCaller (https://github.com/stschiff/sequenceTools/tree/master/src-pileupCaller) for the Mexican specimen with the highest endogenous human content (2179, 2180) at loci that over-lapped with the Affymetrix Human Origins SNP array data (Patterson et al., 2012) and merged them to a modern European, Asian and Native American subset (n=2068) (Lazaridis et al., 2016). Principal Component Analysis (PCA) (Patterson et al., 2006b; Price et al., 2006) on the resulting SNP dataset show that the human DNA form the two coprolite samples has the greatest genetic affinity with modern Native Americans (**Supplementary figure S11**). This result highly supports the haplogroup assignment of the uniparental marker and genetically allocates the specimens to the American continent.

### *P. copri* genome reconstruction from ancient gut metagenomes and molecular dating

To reconstruct ancient *P. copri* genomes, we utilized our in-house scripts (https://bitbucket.org/CibioCM/cmseq) to build 4 ancient *P. copri* genome “scaffolds”, extracting consensus sites of aligned reads of sample 2179 and 2180 (the two samples with all four copri clades detected, Figure 5B) to representative genomes, one for each of the four clades for *P. corpi* complex. Sites covered by ancient reads were filled with gaps if one of following quality criteria was violated: (1) mapping quality is less than 30, (2) coverage is less than 5-fold, (3) the length of aligned read is less than 50nt, (4) minimum identity for the read is less than 97%, (5) minimum dominant allele frequency is less than 80%. Phylogeny of each core gene of a total of 540 was analyzed separately using BEAST v2.5.1 (Bouckaert et al., 2014). Core genes (n=210) supporting monophyly of the 4 *P. copri* clades (thus are unlikely to have been subject of horizontal gene transfer) were kept for searching for orthologs in ancient *P. copri* and their modern counterparts. We searched for these orthologs by aligning selected core genes against 4 “scaffolds” representative of the clades using BLASTn (Altschul et al., 1990) with parameter -word_size of 9. Mapping hits with either length less than 30 bp or e-value over 1e-10 were excluded. We kept orthologs shared by all 72 modern *P. copri* genomes and at least 1 ancient *P. copri* genome “scaffold”, and subsequently applied multiple sequence alignment using mafft (Katoh et al., 2002), with parameter --maxiterate of 1000 and --globalpair, to each of orthologs. Single-ortholog alignments were manually curated excluding mis-aligned sites (we consider continuous variant nucleotides observed in the alignment as artificially mis-aligned sites) and were then merged into one concatenation alignment.

Out of eight ancient strains, we chose only the one with the best overall coverage which was the Clade A strain from sample 2180 (**Supplementary Figure 12**). This sample was accurately radiocarbon dated (A.D. 673 to 768, 1284 ± 16 B.P). The alignment composed of the selected ancient *P. copri* starin and 72 modern strains, which was further processed to automatically remove gappy columns using trimAl(Capella-Gutiérrez et al., 2009). The final alignment included 214,399 nucleotide positions. BEAST v2.5.1(Bouckaert et al., 2014) was used to infer divergence times of *P. copri* clades, using a GTR model of nucleotide substitution (with 4 gamma categories). To choose the best clock and demographic models we performed a model selection comparing coalescent constant, coalescent exponential, coalescent bayesian skyline, and coalescent extended bayesian skyline models (for the demographic priors) and strict and relaxed lognormal (for the clock prior). Model selection (**Supplementary Table S10**) was performed by comparing AICM from BEAST analyses with 100,000,000 Markov Chain Monte Carlo (MCMC) states for each model and sampling every 10,000 states. Convergence of posteriors was assessed by visualising log files with Tracer v1.7(Rambaut et al., 2018). The most fitting combination of models was a coalescent constant population, with strict clock: this analysis was run longer for 204,000,000 iterations and effective sample size (ESS) of all parameters was over 200.

### Data availability

The 15 isolate *P. copri* genomes (and the extended set of 83 isolate, see above) and all metegenomically assembled metagenomes (MAGS) are available here: http://segatalab.cibio.unitn.it/data/Pcopri_Tett_et_al.html and metadata given in **Supplemnetary Tables S2, S4**. The full *P. copri* phylogeny of 1023 genomes is available as **Supplementary Data File 1** (Newick format) at http://segatalab.cibio.unitn.it/data/Pcopri_Tett_et_al.html. Metadata for the three sequenced non-Westernised dataset is given in **Supplementary Tables S11, S12, S13** and is also available as part of the *curatedMetagenomicData* package (Pasolli et al., 2017). The metagenomic reads for these datasets are available under NCBI-SRA BioProject ids; PRJNA529124 (Ghana), PRJNA529400 (Tanzania), PRJNA504891 (Ethiopia). The Data for the ancient metagenomic samples are available from the European Nucleotide Archive under accession no. PRJEB31971.

## Acknowledgements

We thank Levi Waldron and the members of his laboratory for the effort in developing and supporting *curatedMetagenomicData* (Pasolli et al., 2017), and all the members of the Segata laboratory for fruitful discussions and support. We thank the GeNaPi Project, a collaborative study between Emalaikat Foundation, University of Valencia and IATA-CSIC, for data and sample collection from the Ethiopia cohort, and particularly Mari Olcina and Lourdes Larruy. We thank all field workers and laboratory technicians involved in the sample collection in Ghana and Tanzania. This work was supported by NIH NHGRI grant R01HG005220, NIDDK grant R24DK110499, NIDDK grant U54DE023798, CMIT grant 6935956 to C.H., the NYU-HHC CTSI grant (1TL1TR001447) to H.F.P., the U.S. National Institute of Health grant R01-DK103358-01 to R.B. and D.R.L., the Howard Hughes Medical Institute to D.R.L. Further support was provided by the Programma Ricerca Budget prestazioni Eurac 2017 of the Province of Bolzano, Italy to F.M., by the EU-H2020 (DiMeTrack-707345) to E.P. and by the European Research Council (ERC-STG project MetaPG-716575), MIUR ‘‘Futuro in Ricerca’’ RBFR13EWWI_001, and the European Union (H2020-SFS-2018-1 project MASTER-818368) to N.S.

## Supplementary Material

### Supplementary Tables/figures as a single pdf file

**Table S1**, Metagenomic datasets included in this study

**Table S3**, Non-Westernised datasets recently sequenced and sequenced as part of this study

**Figure S1**, Genome statistics for automatically metagenimically assembled genomes (MAGs), manually curated mategenomic assembled genomes (MC-MAGs) and isolate sequences

**Figure S2**, Prevalence and abundance of the *P. copri* complex in publicly available datasets for which there are case and control samples

**Figure S3**, Prevalence of *P. copri* complex in healthy publicly available metagenomes for which Body-Mass-Index (BMI) information was available

**Figure S4**, Prevalence of *P. copri* complex in healthy publicly available metagenomes for which age of participants was available

**Figure S5**, Phylogeny of all 1121 *P. copri* genomes recovered in this study (including 98 new non-Westernised genomes)

**Figure S6**, Prevalence of *P. copri* clades in healthy individuals for datasets considered in this study

**Figure S7**, The within sample *P. copri* complex pangenome for datasets considered in this study

**Figure S8**, Presence absence of CAZy families in each of the 1121 *P. copri* genomes

**Figure S9**, CAZy families present and their prevalence in each clade of the *P. copri* complex

**Figure S10**, DNA damage patterns in ancient metagenomic samples

**Figure S11**, PCA plot of two Mexican coprolite samples and selected modern European, Asian and Native American

**Figure S12**, Depth and breadth of metagenomic reads from sample 2180 mapped to four *P. copri* isolate genomes representing the four *P. copri* complex clades

### Supplementary Tables S2 and S4-12 as a multi-tab Excel file

**Table S2**, List of the 1,023 *P. copri* genomes and their associated genome statistics.

**Table S4**, The list of the reconstructed 98 *P. copri* genomes from the newly sequenced datasets and their associated genome statistics

**Table S5**, Prevalence of eggNOG categories in each of the *P. copri* complex clades

**Table S6**, CAZy prevalence in each of the *P. copri* complex clades

**Table S7**, Description of the ancient metagenomic samples used in this study

**Table S8**, Radiocarbon data for coprolite sample 2180

**Table S9**, Mapping result for the ancient samples versus human and *P. copri* genomes

**Table S10**, Model selection for the BEAST2 phylogeny

**Table S11**, Metadata for the newly sequenced CM_Tanzania metagenomic dataset

**Table S12**, Metadata for the newly sequenced CM_Ghana metagenomic dataset

**Table S13**, Metadata for the newly sequenced CM_Ethiopia metagenomic dataset

