## Supplementary_Tables_S1_S3_Figures_S1_to_S12 for "The *Prevotella copri* complex comprises four distinct clades that are underrepresented in Westernised populations"

| Dataset_name | Samples | Countries (if known) | Reference |
|---|---|---|---|
| AsnicarF_2017 | 16 | ITA | (Asnicar et al., 2017) |
| BackhedF_2015 | 381 | SWE | (Bäckhed et al., 2015) |
| Bengtsson-PalmeJ_2015 | 70 | SWE | (Bengtsson-Palme et al., 2015) |
| BritoIL_2016 | 172 | FJI | (Brito et al., 2016) |
| ChengpingW_2017 | 97 | CHN | (Wen et al., 2017) |
| FerrettiP_2018 | 118 | ITA | (Ferretti et al., 2018) |
| FengQ_2015 | 154 | AUT | (Feng et al., 2015) |
| GeversD_2014 | 50 | NA | (Gevers et al., 2014) |
| HanniganGD_2017 | 82 | CAN,USA | (Hannigan et al., 2018) |
| HeQ_2017 | 122 | NA,CHN | (He et al., 2017) |
| HMP_2012 | 147 | USA | (Human Microbiome Project Consortium, 2012) |
| KarlssonFH_2013 | 145 | FRA,EST,SVK,HUN,SWE, DEU,DNK,ISL,FIN,NOR | (Karlsson et al., 2013) |
| KosticAD_2015 | 124 | NA,EST,FIN | (Kostic et al., 2015) |
| DavidLA_2015 | 49 | BGD | (David et al., 2015) |
| LeChatelierE_2013 | 292 | DNK | (Le Chatelier et al., 2013) |
| LiJ_2014 | 260 | DNK,CHN,ESP | (Li et al., 2014) |
| LiJ_2017 | 196 | CHN | (Li et al., 2017) |
| LiSS_2016 | 55 | NLD | (Li et al., 2016) |
| LiuW_2016 | 110 | MNG | (Liu et al., 2016) |
| LomanNJ_2013 | 43 | DEU | (Loman et al., 2013) |
| LouisS_2016 | 92 | DEU | (Louis et al., 2016) |
| NielsenHB_2014 | 219 | ESP | (Nielsen et al., 2014) |
| Obregon-TitoAJ_2015 | 58 | USA,PER | (Obregon-Tito et al., 2015) |
| QinJ_2012 | 363 | CHN | (Qin et al., 2012) |
| QinN_2014 | 237 | CHN | (Qin et al., 2014) |
| RampelliS_2015 | 38 | TZA,ITA | (Rampelli et al., 2015) |
| RaymondF_2016 | 72 | CAN | (Raymond et al., 2016) |
| SchirmerM_2016 | 471 | NLD | (Schirmer et al., 2016) |
| SmitsSA_2017 | 40 | TZA | (Smits et al., 2017) |
| VatanenT_2016 | 785 | RUS,EST,FIN | (Vatanen et al., 2016) |
| VincentC_2016 | 229 | CAN | (Vincent et al., 2016) |
| VogtmannE_2016 | 110 | USA | (Vogtmann et al., 2016) |
| XieH_2016 | 250 | GBR | (Xie et al., 2016) |
| YuJ_2015 | 128 | CHN | (Yu et al., 2017) |
| ZeeviD_2015 | 900 | ISR | (Zeevi et al., 2015) |
| ZellerG_2014 | 199 | FRA,DEU | (Zeller et al., 2014) |
| **Total** | **6,874** | | |

**Supplementary Table 1** Publicly available metagenomic datasets considered as part of this study.

| Dataset_name | Samples | Countries | Reference |
|---|---|---|---|
| CM_Ethiopia | 50 | ETH | *This study* |
| CM_Ghana | 44 | GHA | *This study* |
| CM_Madagascar | 110 | MDG | (Pasolli et al., 2019) |
| CM_Tanzania | 68 | TZA | *This study* |

**Supplementary Table 3** Non-Westernised datasets recently sequenced and sequenced as part of this study.

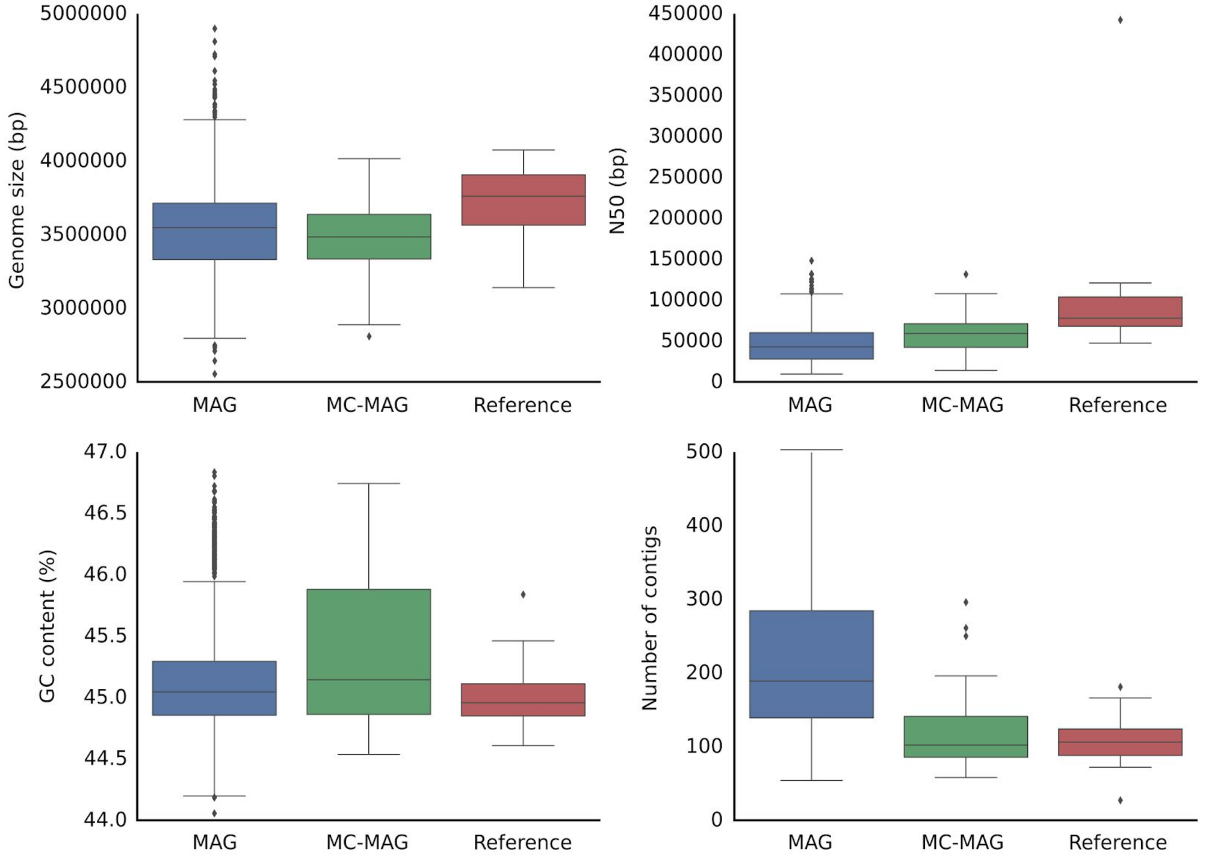


**Supplementary Figure 1.** Comparison of genome statistics (Genome size, N50, GC% and number of contigs) for *P. copri* isolate sequences (references = 17), for manually curated mategenomic assembled genomes (MC-MAG = 55) and automatically metagenomically assembled genomes (MAGs = 951).

| # sample | Prevalence (Fisher) | | | | | Abundance (Mann-Whitney) | | | | | | |
|---|---|---|---|---|---|---|---|---|---|---|---|---|
| | Any | Clade A | Clade B | Clade C | Clade D | Any | Clade A | Clade B | Clade C | Clade D | Dataset | Condition |
| 66 | 27.3 | 21.2 | 7.6 | 12.1 | 3.0 | 6.7 | 4.1 | 0.4 | 7.6 | 0.6 | ZellerG_2014 | Control |
| 42 | 35.7 | 33.3 | 11.9 | 9.5 | 4.8 | 11.6 | 8.8 | 1.5 | 4.2 | 13.5 | | Adenoma |
| 91 | 26.4 | 20.9 | 11.0 | 7.7 | 4.4 | 6.6 | 5.4 | 2.4 | 0.9 | 6.4 | | CRC |
| 133 | 29.3 | 24.8 | 11.3 | 8.3 | 4.5 | 8.6 | 6.8 | 2.1 | 2.1 | 8.8 | | Adenoma/CRC |
| 61 | 8.2 | 3.3 | 1.6 | 3.3 | 1.6 | 1.0 | 0.5 | 1.5 | 0.8 | 1.1 | FengQ_2015 | Control |
| 46 | **39.1** * | **39.1** * | 4.3 | **17.4** * | 0.0 | **7.0** * | **6.4** * | 0.9 | **1.1** * | 0.0 | | CRC |
| 47 | 17.0 | 12.8 | 4.3 | 6.4 | 0.0 | 2.0 | 1.7 | 0.5 | 1.6 | 0.0 | | Adenoma |
| 93 | **28.0** * | **25.8** * | 4.3 | 11.8 | 0.0 | **5.5** * | **5.3** * | 0.7 | 1.2 | 0.0 | | CRC/adenoma |
| 52 | 23.1 | 23.1 | 1.9 | 0.0 | 0.0 | 11.7 | 11.7 | 0.2 | 0.0 | 0.0 | VogtmannE_2016 | Control |
| 52 | 19.2 | 19.2 | 0.0 | 0.0 | 0.0 | 12.6 | 12.6 | 0.0 | 0.0 | 0.0 | | CRC |
| 53 | 28.3 | 24.5 | 1.9 | 7.5 | 0.0 | 10.2 | 9.8 | 0.3 | 6.3 | 0.0 | YuJ_2015 | Control |
| 75 | 30.7 | 28.0 | 5.3 | 4.0 | 0.0 | 10.0 | 10.2 | 2.2 | 2.7 | 0.0 | | CRC |
| 174 | 29.9 | 25.9 | 4.0 | 8.0 | 1.7 | 28.6 | 26.4 | 10.2 | 15.1 | 5.2 | QinJ_2012 | Control |
| 170 | 28.2 | 25.3 | 6.5 | 10.0 | 1.8 | 17.2 | 13.7 | 7.3 | 8.1 | 6.9 | | T2D |
| 43 | 14.0 | 11.6 | 0.0 | 7.0 | 0.0 | 2.1 | 1.6 | 0.0 | 1.5 | 0.0 | KarlssonFH_2013 | Control |
| 102 | 16.7 | 14.7 | 2.0 | 2.9 | 2.0 | 4.4 | 3.7 | 0.5 | 4.0 | 2.8 | | IGT/T2D |
| 49 | 10.2 | 6.1 | 2.0 | 4.1 | 0.0 | 2.6 | 1.5 | 0.2 | 4.1 | 0.0 | | IGT |
| 53 | 22.6 | 22.6 | 1.9 | 1.9 | 3.8 | 5.1 | 4.3 | 0.8 | 4.0 | 2.8 | | T2D |
| 41 | 39.0 | 39.0 | 7.3 | 4.9 | 0.0 | 31.2 | 29.7 | 2.5 | 8.5 | 0.0 | LiJ_2017 | Control |
| 99 | 48.5 | 47.5 | 11.1 | 16.2 | 2.0 | 37.3 | 31.6 | 4.2 | 13.8 | 18.6 | | Hypertension |
| 56 | 53.6 | 48.2 | 3.6 | **19.6** * | 1.8 | 41.4 | 39.0 | 11.4 | **14.9** * | 2.1 | | Pre-hypertension |
| 155 | 50.3 | 47.7 | 8.4 | **17.4** * | 1.9 | 38.9 | 34.3 | 5.3 | **14.2** * | 13.1 | | Pre-hypertension/hypertension |
| 71 | 25.4 | 25.4 | 1.4 | 2.8 | 0.0 | 8.6 | 8.1 | 0.4 | 4.6 | 0.0 | NielsenHB_2014 | Control |
| 148 | 33.8 | 29.7 | **11.5** * | 3.4 | **6.1** * | 11.5 | 10.1 | **2.8** * | 8.9 | **4.5** * | | IBD |
| 53 | 26.4 | 24.5 | 1.9 | 11.3 | 3.8 | 15.1 | 12.5 | 1.1 | 6.7 | 3.4 | HeQ_2017 | Control |
| 63 | 12.7 | 11.1 | 4.8 | 3.2 | 0.0 | 45.9 | 31.6 | 17.6 | 46.6 | 0.0 | | CD |
| 114 | 52.6 | 48.2 | 1.8 | 13.2 | 1.8 | 9.6 | 9.1 | 4.8 | 3.9 | 2.1 | QinN_2014 | Control |
| 123 | 58.5 | 57.7 | 2.4 | 16.3 | 0.8 | 11.7 | 9.7 | 1.5 | 6.8 | 14.5 | | Cirrhosis |
| 36 | 11.1 | 8.3 | 5.6 | 2.8 | 0.0 | 25.4 | 26.2 | 10.0 | 3.0 | 0.0 | RaymondF_2016 | Control |
| 36 | 19.4 | 13.9 | 11.1 | 5.6 | 0.0 | 16.0 | 17.6 | 4.6 | 2.8 | 0.0 | | Cephalosporins |

**Supplementary Figure 2.** Prevalence and abundance of the *P. copri* complex in publicly available datasets for which there are case and control samples. ***** indicates p <0.05 (Fisher exact test (prevalence) or Mann-Whitney (abundance))

| Dataset | BMI | # Samples | Prevalence Any | Clade A | Clade B | Clade C | Clade D |
|---|---|---|---|---|---|---|---|
| NielsenHB_2014 | UNDER | 2 | 0.0 | 0.0 | 0.0 | 0.0 | 0.0 |
| | NORMAL | 35 | 37.1 | 37.1 | 0.0 | 5.7 | 0.0 |
| | OVER | 17 | 11.8 | 11.8 | 5.9 | 0.0 | 0.0 |
| | OBESE | 5 | 20.0 | 20.0 | 0.0 | 0.0 | 0.0 |
| QinN_2014 | UNDER | 2 | 0.0 | 0.0 | 0.0 | 0.0 | 0.0 |
| | NORMAL | 105 | 52.4 | 48.6 | 1.9 | 12.4 | 1.0 |
| | OVER | 7 | 71.4 | 57.1 | 0.0 | 28.6 | 14.3 |
| XieH_2016 | UNDER | 4 | 75.0 | 75.0 | 25.0 | 25.0 | 25.0 |
| | NORMAL | 129 | 21.7 | 21.7 | 2.3 | 1.6 | 0.8 |
| | OVER | 71 | 18.3 | 16.9 | 1.4 | 9.9 | 2.8 |
| | OBESE | 45 | 8.9 | 8.9 | 0.0 | 0.0 | 0.0 |
| LeChatelierE_2013 | UNDER | 1 | 100.0 | 100.0 | 0.0 | 100.0 | 0.0 |
| | NORMAL | 94 | 35.1 | 31.9 | 5.3 | 3.2 | 4.3 |
| | OVER | 28 | 35.7 | 35.7 | 10.7 | 7.1 | 0.0 |
| | OBESE | 169 | 37.9 | 32.5 | 10.1 | 8.3 | 5.9 |
| RaymondF_2016 | NORMAL | 31 | 6.5 | 3.2 | 3.2 | 0.0 | 0.0 |
| | OVER | 5 | 40.0 | 40.0 | 20.0 | 20.0 | 0.0 |
| | OBESE | 0 | 0.0 | 0.0 | 0.0 | 0.0 | 0.0 |
| HeQ_2017 | UNDER | 5 | 20.0 | 20.0 | 0.0 | 0.0 | 0.0 |
| | NORMAL | 40 | 27.5 | 25.0 | 2.5 | 12.5 | 5.0 |
| | OVER | 7 | 28.6 | 28.6 | 0.0 | 14.3 | 0.0 |
| | OBESE | 1 | 0.0 | 0.0 | 0.0 | 0.0 | 0.0 |
| FengQ_2015 | NORMAL | 20 | 5.0 | 5.0 | 0.0 | 0.0 | 0.0 |
| | OVER | 20 | 15.0 | 0.0 | 5.0 | 10.0 | 5.0 |
| | OBESE | 21 | 4.8 | 4.8 | 0.0 | 0.0 | 0.0 |
| Obregon-TitoAJ_2015 | UNDER | 13 | 76.9 | 61.5 | 61.5 | 69.2 | 61.5 |
| | NORMAL | 26 | 57.7 | 53.8 | 42.3 | 46.2 | 26.9 |
| | OVER | 9 | 55.6 | 55.6 | 33.3 | 33.3 | 33.3 |
| | OBESE | 3 | 33.3 | 33.3 | 33.3 | 33.3 | 33.3 |
| VogtmannE_2016 | UNDER | 1 | 0.0 | 0.0 | 0.0 | 0.0 | 0.0 |
| | NORMAL | 31 | 25.8 | 25.8 | 3.2 | 0.0 | 0.0 |
| | OVER | 12 | 16.7 | 16.7 | 0.0 | 0.0 | 0.0 |
| | OBESE | 8 | 25.0 | 25.0 | 0.0 | 0.0 | 0.0 |
| QinJ_2012 | UNDER | 17 | 52.9 | 47.1 | 5.9 | 5.9 | 0.0 |
| | NORMAL | 96 | 26.0 | 21.9 | 5.2 | 7.3 | 2.1 |
| | OVER | 59 | 28.8 | 25.4 | 1.7 | 10.2 | 1.7 |
| | OBESE | 2 | 50.0 | 50.0 | 0.0 | 0.0 | 0.0 |
| ZellerG_2014 | UNDER | 1 | 100.0 | 100.0 | 100.0 | 100.0 | 0.0 |
| | NORMAL | 31 | 19.4 | 16.1 | 3.2 | 9.7 | 0.0 |
| | OVER | 25 | 28.0 | 20.0 | 8.0 | 8.0 | 8.0 |
| | OBESE | 7 | 42.9 | 28.6 | 14.3 | 28.6 | 0.0 |
| FerrettiP_2018 | UNDER | 4 | 0.0 | 0.0 | 0.0 | 0.0 | 0.0 |
| | NORMAL | 15 | 20.0 | 20.0 | 13.3 | 6.7 | 6.7 |
| | OVER | 3 | 33.3 | 33.3 | 0.0 | 0.0 | 0.0 |
| SchirmerM_2016 | UNDER | 14 | 28.6 | 28.6 | 14.3 | 0.0 | 0.0 |
| | NORMAL | 366 | 39.9 | 36.1 | 7.1 | 6.6 | 4.1 |
| | OVER | 68 | 50.0 | 47.1 | 10.3 | 11.8 | 5.9 |
| | OBESE | 8 | 37.5 | 37.5 | 12.5 | 0.0 | 0.0 |
| **Total** | **UNDER** | **64** | **45.3** | **40.6** | **20.3** | **20.3** | **14.1** |
| | **NORMAL** | **1019** | **34.0** | **31.1** | **5.7** | **7.1** | **3.2** |
| | **OVER** | **331** | **31.1** | **27.8** | **6.0** | **10.3** | **4.2** |
| | **OBESE** | **269** | **29.7** | **26.0** | **7.4** | **6.3** | **4.1** |

**Supplementary Figure 3.** Prevalence of *P. copri* complex in healthy publicly available metagenomes for which Body-Mass-Index (BMI) information was available (Under <18.5, Normal ≥18.5 <25, Overweight ≥25 <30, Obese ≥30).

| Dataset | AGE | #Samples | Prevalence | | | | |
|---|---|---|---|---|---|---|---|
| | | | Any | Clade A | Clade B | Clade C | Clade D |
| NielsenHB_2014 | Adult | 66 | 25.8 | 25.8 | 1.5 | 3.0 | 0.0 |
| | Elderly | 4 | 0.0 | 0.0 | 0.0 | 0.0 | 0.0 |
| FengQ_2015 | Adult | 14 | 7.1 | 7.1 | 0.0 | 0.0 | 0.0 |
| | Elderly | 47 | 8.5 | 2.1 | 2.1 | 4.3 | 2.1 |
| Obregon-TitoAJ_2015 | Infant | 5 | 100.0 | 100.0 | 100.0 | 100.0 | 40.0 |
| | School_age | 16 | 81.3 | 68.8 | 62.5 | 68.8 | 68.8 |
| | Adult | 36 | 52.8 | 50.0 | 36.1 | 36.1 | 30.6 |
| VatanenT_2016 | Infant | 616 | 13.0 | 12.3 | 2.1 | 2.1 | 0.0 |
| FerrettiP_2018 | Infant | 96 | 5.2 | 5.2 | 0.0 | 2.1 | 0.0 |
| | Adult | 19 | 10.5 | 10.5 | 5.3 | 5.3 | 5.3 |
| QinN_2014 | Adult | 112 | 53.6 | 49.1 | 1.8 | 13.4 | 1.8 |
| RampelliS_2015 | School_age | 5 | 100.0 | 100.0 | 80.0 | 100.0 | 40.0 |
| | Adult | 31 | 74.2 | 67.7 | 61.3 | 64.5 | 35.5 |
| | Elderly | 2 | 100.0 | 100.0 | 100.0 | 100.0 | 100.0 |
| BritoIL_2016 | Infant | 11 | 100.0 | 100.0 | 100.0 | 100.0 | 90.9 |
| | School_age | 42 | 100.0 | 100.0 | 83.3 | 100.0 | 76.2 |
| | Adult | 114 | 100.0 | 100.0 | 91.2 | 100.0 | 91.2 |
| | Elderly | 5 | 100.0 | 100.0 | 80.0 | 100.0 | 60.0 |
| XieH_2016 | Adult | 153 | 16.3 | 15.7 | 2.6 | 2.6 | 0.7 |
| | Elderly | 97 | 23.7 | 23.7 | 1.0 | 6.2 | 3.1 |
| RaymondF_2016 | Adult | 36 | 11.1 | 8.3 | 5.6 | 2.8 | 0.0 |
| HeQ_2017 | School_age | 26 | 23.1 | 23.1 | 0.0 | 7.7 | 0.0 |
| | Adult | 27 | 29.6 | 25.9 | 3.7 | 14.8 | 7.4 |
| VincentC_2016 | Adult | 3 | 0.0 | 0.0 | 0.0 | 0.0 | 0.0 |
| | Elderly | 24 | 8.3 | 4.2 | 4.2 | 4.2 | 0.0 |
| AsnicarF_2017 | Infant | 8 | 12.5 | 12.5 | 0.0 | 0.0 | 0.0 |
| QinJ_2012 | School_age | 1 | 0.0 | 0.0 | 0.0 | 0.0 | 0.0 |
| | Adult | 167 | 29.3 | 25.7 | 3.6 | 8.4 | 1.8 |
| | Elderly | 6 | 50.0 | 33.3 | 16.7 | 0.0 | 0.0 |
| LeChatelierE_2013 | Adult | 157 | 33.1 | 29.9 | 7.0 | 7.6 | 5.7 |
| | Elderly | 20 | 20.0 | 20.0 | 0.0 | 0.0 | 0.0 |
| ZellerG_2014 | Adult | 43 | 25.6 | 23.3 | 7.0 | 11.6 | 2.3 |
| | Elderly | 23 | 30.4 | 17.4 | 8.7 | 13.0 | 4.3 |
| SmitsSA_2017 | School_age | 8 | 75.0 | 62.5 | 12.5 | 62.5 | 50.0 |
| | Adult | 16 | 93.8 | 81.3 | 25.0 | 75.0 | 43.8 |
| VogtmannE_2016 | Adult | 31 | 29.0 | 29.0 | 3.2 | 0.0 | 0.0 |
| | Elderly | 21 | 14.3 | 14.3 | 0.0 | 0.0 | 0.0 |
| HMP_2012 | Adult | 147 | 15.6 | 15.6 | 3.4 | 4.1 | 0.7 |
| KarlssonFH_2013 | Elderly | 43 | 14.0 | 11.6 | 0.0 | 7.0 | 0.0 |
| Bengtsson-PalmeJ_2015 | Adult | 70 | 40.0 | 34.3 | 2.9 | 14.3 | 2.9 |
| SchirmerM_2016 | Adult | 450 | 41.1 | 37.6 | 7.8 | 6.9 | 4.0 |
| | Elderly | 15 | 40.0 | 40.0 | 6.7 | 13.3 | 6.7 |
| **All datasets** | **Infant** | **736** | **13.9** | **13.3** | **3.9** | **4.2** | **1.6** |
| | **School_age** | **98** | **73.5** | **70.4** | **51.0** | **66.3** | **50.0** |
| | **Adult** | **1692** | **38.1** | **35.5** | **12.6** | **15.6** | **10.2** |
| | **Elderly** | **309** | **21.0** | **18.1** | **4.2** | **7.8** | **3.6** |

**Supplementary Figure 4.** Prevalence of *P. copri* complex in healthy publicly available metagenomes for which age of participants was available. (Infant < 4 yrs; 4 yrs ≤ School age < 18 yrs; 18 yrs ≤ Adult < 65 yrs; Elderly ≥65).

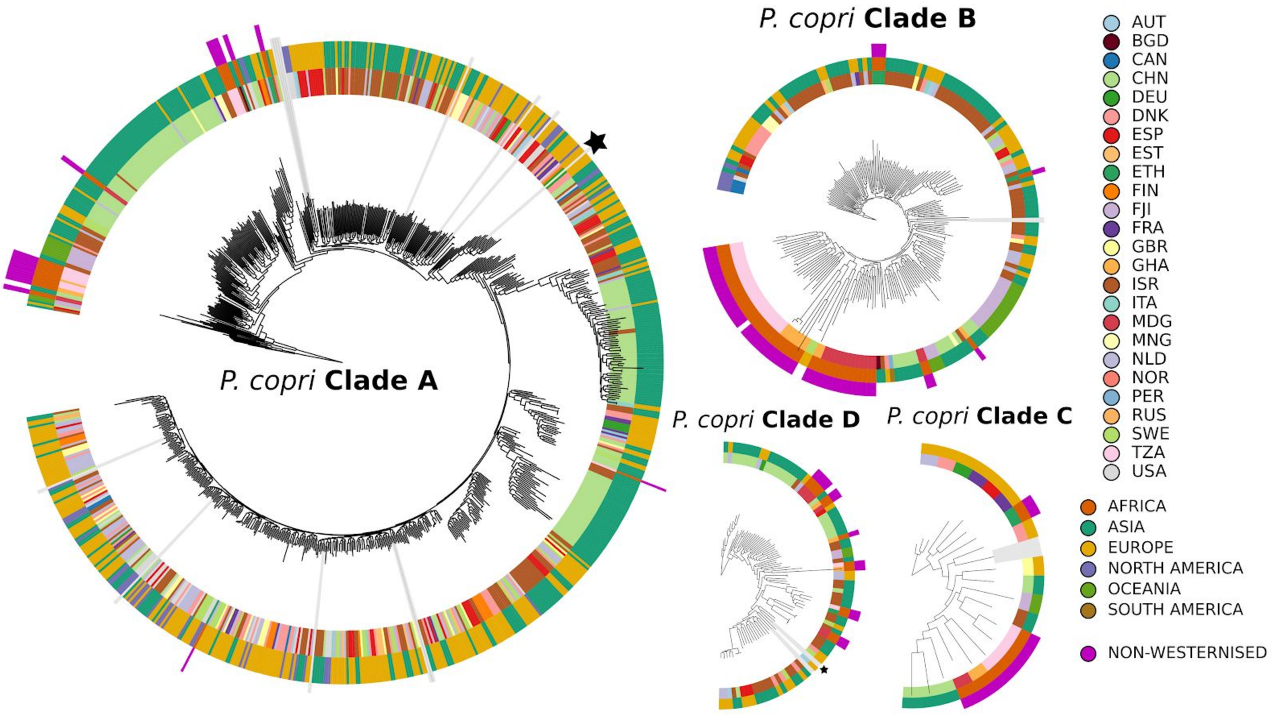


**Supplementary Figure 5.** Phylogeny of all 1121 *P. copri* genomes recovered in this study (including 98 new non-Westernised genomes). Outermost ring indicates if the genome was recovered from a non-Westernised sample, middle ring the continent and inner ring the country of origin. Publicly available *P. copri* references are indicated by black stars and newly isolated genomes by radial grey bars.

| Dataset | Country | #Samples | Any | Clade A | Clade B | Clade C | Clade D |
|---|---|---|---|---|---|---|---|
| | | | Prevalence | | | | |
| FengQ_2015 | AUT | 61 | 8.2 | 3.3 | 1.6 | 3.3 | 1.6 |
| RaymondF_2016 | CAN | 36 | 11.1 | 8.3 | 5.6 | 2.8 | 0.0 |
| VincentC_2016 | CAN | 25 | 8.0 | 4.0 | 4.0 | 4.0 | 0.0 |
| HeQ_2017 | CHN | 53 | 26.4 | 24.5 | 1.9 | 11.3 | 3.8 |
| LiJ_2014 | CHN | 11 | 27.3 | 18.2 | 9.1 | 9.1 | 0.0 |
| LiJ_2017 | CHN | 41 | 39.0 | 39.0 | 7.3 | 4.9 | 0.0 |
| QinJ_2012 | CHN | 174 | 29.9 | 25.9 | 4.0 | 8.0 | 1.7 |
| QinN_2014 | CHN | 114 | 52.6 | 48.2 | 1.8 | 13.2 | 1.8 |
| YuJ_2015 | CHN | 53 | 28.3 | 24.5 | 1.9 | 7.5 | 0.0 |
| LouisS_2016 | DEU | 92 | 12.0 | 12.0 | 0.0 | 0.0 | 0.0 |
| LeChatelierE_2013 | DNK | 292 | 37.0 | 32.9 | 8.6 | 6.8 | 4.8 |
| LiJ_2014 | DNK | 109 | 33.9 | 32.1 | 7.3 | 9.2 | 6.4 |
| NielsenHB_2014 | ESP | 71 | 25.4 | 25.4 | 1.4 | 2.8 | 0.0 |
| KosticAD_2015 | EST | 99 | 2.0 | 2.0 | 1.0 | 0.0 | 0.0 |
| VatanenT_2016 | EST | 156 | 20.5 | 17.9 | 3.8 | 5.8 | 0.0 |
| KosticAD_2015 | FIN | 21 | 28.6 | 28.6 | 14.3 | 0.0 | 0.0 |
| VatanenT_2016 | FIN | 179 | 8.4 | 8.4 | 1.1 | 0.0 | 0.0 |
| ZellerG_2014 | FRA | 61 | 27.9 | 23.0 | 8.2 | 11.5 | 3.3 |
| XieH_2016 | GBR | 250 | 19.2 | 18.8 | 2.0 | 4.0 | 1.6 |
| ZeeviD_2015 | ISR | 900 | 51.4 | 46.1 | 12.7 | 18.1 | 4.4 |
| AsnicarF_2017 | ITA | 16 | 37.5 | 37.5 | 0.0 | 6.3 | 0.0 |
| FerrettiP_2018 | ITA | 118 | 7.6 | 7.6 | 1.7 | 2.5 | 0.8 |
| RampelliS_2015 | ITA | 11 | 27.3 | 18.2 | 9.1 | 0.0 | 0.0 |
| LiuW_2016 | MNG | 45 | 100.0 | 100.0 | 73.3 | 80.0 | 42.2 |
| SchirmerM_2016 | NLD | 471 | 41.0 | 37.4 | 8.1 | 7.0 | 4.0 |
| VatanenT_2016 | RUS | 279 | 11.5 | 11.5 | 1.4 | 1.4 | 0.0 |
| BackhedF_2015 | SWE | 377 | 8.2 | 7.4 | 0.8 | 1.3 | 0.3 |
| Bengtsson-PalmeJ_2015 | SWE | 70 | 40.0 | 34.3 | 2.9 | 14.3 | 2.9 |
| KarlssonFH_2013 | SWE | 39 | 15.4 | 12.8 | 0.0 | 7.7 | 0.0 |
| HanniganGD_2017 | USA | 25 | 4.0 | 4.0 | 0.0 | 0.0 | 0.0 |
| HMP_2012 | USA | 147 | 15.6 | 15.6 | 3.4 | 4.1 | 0.7 |
| Obregon-TitoAJ_2015 | USA | 22 | 13.6 | 13.6 | 4.5 | 9.1 | 0.0 |
| VogtmannE_2016 | USA | 52 | 23.1 | 23.1 | 1.9 | 0.0 | 0.0 |
| BritoIL_2016 | FJI | 172* | 100.0 | 100.0 | 89.5 | 100.0 | 86.6 |
| LiuW_2016 | MNG | 65* | 98.5 | 98.5 | 76.9 | 84.6 | 56.9 |
| Obregon-TitoAJ_2015 | PER | 36* | 97.2 | 88.9 | 77.8 | 77.8 | 69.4 |
| RampelliS_2015 | TZA | 27* | 100.0 | 96.3 | 88.9 | 100.0 | 55.6 |
| SmitsSA_2017 | TZA | 40* | 85.0 | 75.0 | 35.0 | 70.0 | 42.5 |
| CM_Ethiopia | ETH | 50* | 94.0 | 92.0 | 68.0 | 90.0 | 64.0 |
| CM_Ghana | GHA | 44* | 90.9 | 77.3 | 54.5 | 86.4 | 54.5 |
| CM_Madagascar | MDG | 110* | 95.5 | 89.1 | 70.9 | 89.1 | 77.3 |
| CM_Tanzania | TZA | 68* | 88.2 | 85.3 | 64.7 | 72.1 | 54.4 |

**Supplementary Figure S6.** Prevalence of *P. copri* clades in healthy individuals for each dataset. When a dataset included multiple countries they were further divided based on the country of origin (If >10 metagenomes from any given country was available, if not data not shown). * Denotes non-Westernised samples.

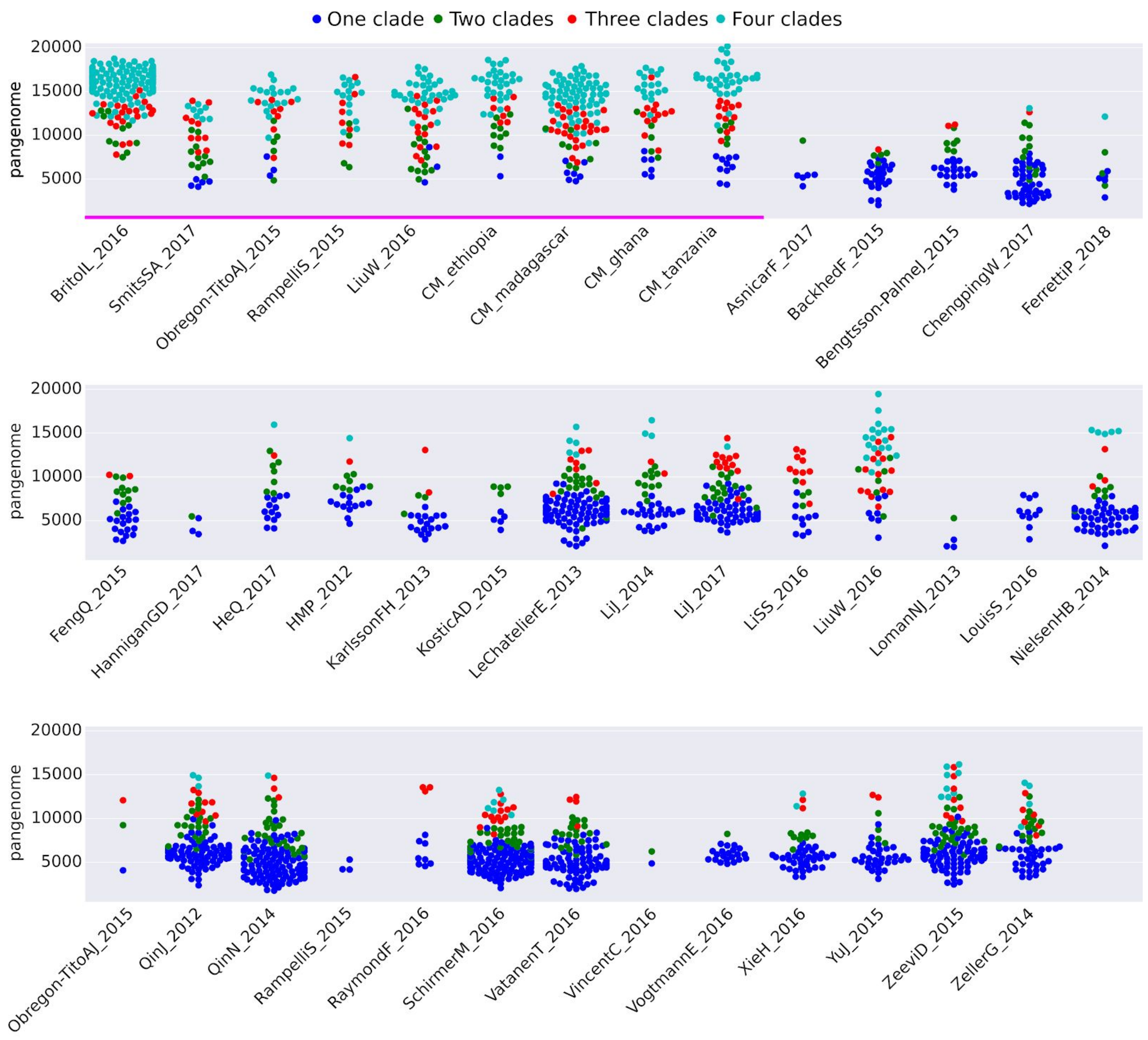


**Supplementary Figure S7.** The within sample *P. copri* complex pangenome for datasets considered in this study. Non-Westernised datasets are underlined in magenta.

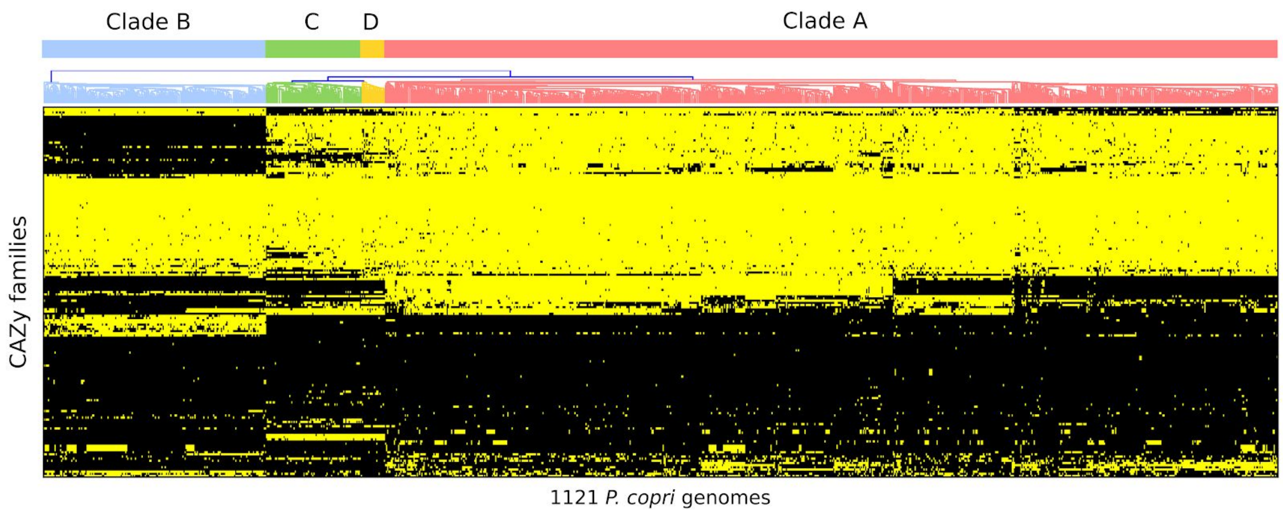


**Supplementary Figure 8.** Presence absence of CAZy families in each of the 1121 *P. copri* genomes (yellow present, black absent)

| | Cazy_family | Clade A | Clade B | Clade C | Clade D | Annotation |
|---|---|---|---|---|---|---|
| Glycoside Hydrolase (GH) | GH8.hmm | 80.15 | 0.00 | 56.32 | 71.43 | Cellulose |
| | GH9.hmm | 2.10 | 0.00 | 33.33 | 80.95 | Cellulose |
| | GH51.hmm | 100.00 | 0.50 | 100.00 | 100.00 | Cellulose/Hemicellulose |
| | GH10.hmm | 100.00 | 0.00 | 98.85 | 100.00 | Hemicellulose |
| | GH115.hmm | 99.63 | 93.56 | 97.70 | 100.00 | Hemicellulose |
| | GH16.hmm | 86.93 | 90.10 | 68.97 | 38.10 | Hemicellulose |
| | GH26.hmm | 96.67 | 3.96 | 75.86 | 80.95 | Hemicellulose |
| | GH27.hmm | 0.25 | 0.00 | 86.21 | 80.95 | Hemicellulose |
| | GH29.hmm | 72.87 | 0.50 | 100.00 | 100.00 | Hemicellulose |
| | GH30_2.hmm | 0.00 | 0.00 | 11.49 | 9.52 | Hemicellulose |
| | GH30_4.hmm | 47.60 | 0.00 | 42.53 | 0.00 | Hemicellulose |
| | GH30.hmm | 69.79 | 91.09 | 0.00 | 0.00 | Hemicellulose |
| | GH31.hmm | 99.63 | 99.50 | 100.00 | 100.00 | Hemicellulose |
| | GH36.hmm | 99.75 | 79.21 | 97.70 | 90.48 | Hemicellulose |
| | GH67.hmm | 97.04 | 0.00 | 97.70 | 95.24 | Hemicellulose |
| | GH95.hmm | 100.00 | 100.00 | 100.00 | 100.00 | Hemicellulose |
| | GH105.hmm | 99.01 | 100.00 | 60.92 | 100.00 | Pectin |
| | GH106.hmm | 71.27 | 100.00 | 41.38 | 90.48 | Pectin |
| | GH138.hmm | 56.23 | 0.00 | 0.00 | 0.00 | Pectin |
| | GH140.hmm | 57.09 | 73.27 | 0.00 | 0.00 | Pectin |
| | GH141.hmm | 53.02 | 0.00 | 0.00 | 0.00 | Pectin |
| | GH142.hmm | 56.97 | 0.00 | 0.00 | 0.00 | Pectin |
| | GH143.hmm | 56.97 | 0.00 | 0.00 | 0.00 | Pectin |
| | GH28.hmm | 99.51 | 100.00 | 96.55 | 100.00 | Pectin |
| | GH53.hmm | 98.77 | 27.72 | 97.70 | 90.48 | Pectin |
| | GH78.hmm | 70.04 | 98.02 | 9.20 | 33.33 | Pectin |
| | GH2.hmm | 100.00 | 99.50 | 100.00 | 100.00 | Pectin also mucin |
| | GH35.hmm | 75.09 | 0.00 | 70.11 | 100.00 | Pectin/Hemicellulose |
| | **GH43_family** | | | | | Pectin/Hemicellulose |
| | GH43_1.hmm | 98.64 | 0.00 | 96.55 | 80.95 | |
| | GH43_10.hmm | 100.00 | 99.50 | 97.70 | 100.00 | |
| | GH43_12.hmm | 98.40 | 0.50 | 90.80 | 100.00 | |
| | GH43_17.hmm | 0.25 | 0.00 | 16.09 | 0.00 | |
| | GH43_18.hmm | 56.35 | 0.00 | 0.00 | 0.00 | |
| | GH43_19.hmm | 60.79 | 0.00 | 71.26 | 90.48 | |
| | GH43_2.hmm | 75.83 | 0.00 | 68.97 | 76.19 | |
| | GH43_24.hmm | 39.83 | 0.50 | 49.43 | 95.24 | |
| | GH43_28.hmm | 0.00 | 52.97 | 0.00 | 0.00 | |
| | GH43_29.hmm | 98.03 | 0.00 | 89.66 | 80.95 | |
| | GH43_3.hmm | 9.86 | 0.00 | 0.00 | 0.00 | |
| | GH43_31.hmm | 10.36 | 0.00 | 0.00 | 0.00 | |
| | GH43_35.hmm | 11.96 | 0.00 | 1.15 | 4.76 | |
| | GH43_4.hmm | 98.03 | 0.00 | 96.55 | 85.71 | |
| | GH43_5.hmm | 97.04 | 0.00 | 95.40 | 85.71 | |
| | GH43_7.hmm | 89.52 | 0.00 | 74.71 | 80.95 | |
| | GH43_9.hmm | 0.49 | 0.00 | 0.00 | 14.29 | |
| | GH43.hmm | 93.71 | 0.50 | 55.17 | 90.48 | |
| | GH3.hmm | 100.00 | 100.00 | 100.00 | 100.00 | Pectin/Hemicellulose/cellulose |
| | **GH5_family** | | | | | Pectin/Hemicellulose/cellulose |
| | GH5_2.hmm | 71.89 | 12.38 | 0.00 | 90.48 | |
| | GH5_21.hmm | 95.44 | 0.00 | 64.37 | 80.95 | |
| | GH5_25.hmm | 14.30 | 15.35 | 2.30 | 9.52 | |
| | GH5_35.hmm | 0.00 | 1.98 | 0.00 | 0.00 | |
| | GH5_4.hmm | 99.75 | 19.80 | 26.44 | 80.95 | |
| | GH5_46.hmm | 1.36 | 0.00 | 6.90 | 9.52 | |
| | GH5_5.hmm | 0.00 | 13.86 | 0.00 | 0.00 | |
| | GH5_7.hmm | 95.07 | 3.96 | 5.75 | 66.67 | |
| | GH5.hmm | 0.25 | 0.00 | 0.00 | 0.00 | |
| | **GH13_family** | | | | | Starch |
| | GH13_13.hmm | 98.03 | 100.00 | 96.55 | 90.48 | |
| | GH13_19.hmm | 5.55 | 0.00 | 6.90 | 0.00 | |
| | GH13_2.hmm | 0.25 | 0.00 | 0.00 | 0.00 | |
| | GH13_20.hmm | 0.00 | 19.31 | 3.45 | 0.00 | |
| | GH13_36.hmm | 84.46 | 43.07 | 78.16 | 80.95 | |
| | GH13_38.hmm | 95.07 | 98.51 | 89.66 | 95.24 | |
| | GH13_42.hmm | 3.21 | 7.43 | 0.00 | 0.00 | |
| | GH13_6.hmm | 0.49 | 0.99 | 0.00 | 0.00 | |
| | GH13_7.hmm | 96.67 | 94.55 | 97.70 | 85.71 | |
| | GH13_8.hmm | 98.89 | 99.50 | 100.00 | 100.00 | |
| | GH13.hmm | 98.64 | 100.00 | 98.85 | 90.48 | |
| | GH57.hmm | 99.26 | 97.52 | 100.00 | 95.24 | Starch |
| | GH63.hmm | 48.58 | 0.00 | 0.00 | 0.00 | Starch |
| | GH77.hmm | 99.14 | 99.01 | 97.70 | 95.24 | Starch |
| | GH97.hmm | 100.00 | 100.00 | 100.00 | 100.00 | Starch |
| Carbohydrate Esterases (GE) | CE1.hmm | 100.00 | 100.00 | 100.00 | 100.00 | Hemicellulose |
| | CE2.hmm | 95.81 | 1.98 | 1.15 | 90.48 | Hemicellulose |
| | CE3.hmm | 0.00 | 68.81 | 0.00 | 0.00 | Hemicellulose |
| | CE4.hmm | 99.63 | 99.01 | 97.70 | 100.00 | Hemicellulose |
| | CE6.hmm | 96.30 | 0.00 | 82.76 | 90.48 | Hemicellulose |
| | CE7.hmm | 97.90 | 99.50 | 73.56 | 100.00 | Hemicellulose |
| | CE8.hmm | 100.00 | 100.00 | 54.02 | 100.00 | Pectin |
| | CE12.hmm | 99.26 | 99.01 | 49.43 | 100.00 | Pectin/Hemicellulose |
| Polysaccharide Lyases (PL) | PL1_2.hmm | 87.92 | 98.02 | 8.05 | 66.67 | Pectin |
| | PL1.hmm | 54.62 | 91.58 | 0.00 | 52.38 | Pectin |
| | PL10.hmm | 2.10 | 98.02 | 0.00 | 0.00 | Pectin |
| | PL11.hmm | 34.65 | 0.50 | 33.33 | 90.48 | Pectin |
| | PL4.hmm | 0.00 | 0.99 | 0.00 | 0.00 | Pectin |
| | PL9.hmm | 0.86 | 97.03 | 3.45 | 0.00 | Pectin |

**Supplementary Figure 9.** CAZy families present and their prevalence in each clade of the *P. copri* complex. Families are separated based on their enzyme class and broad predicted substrate specificity.

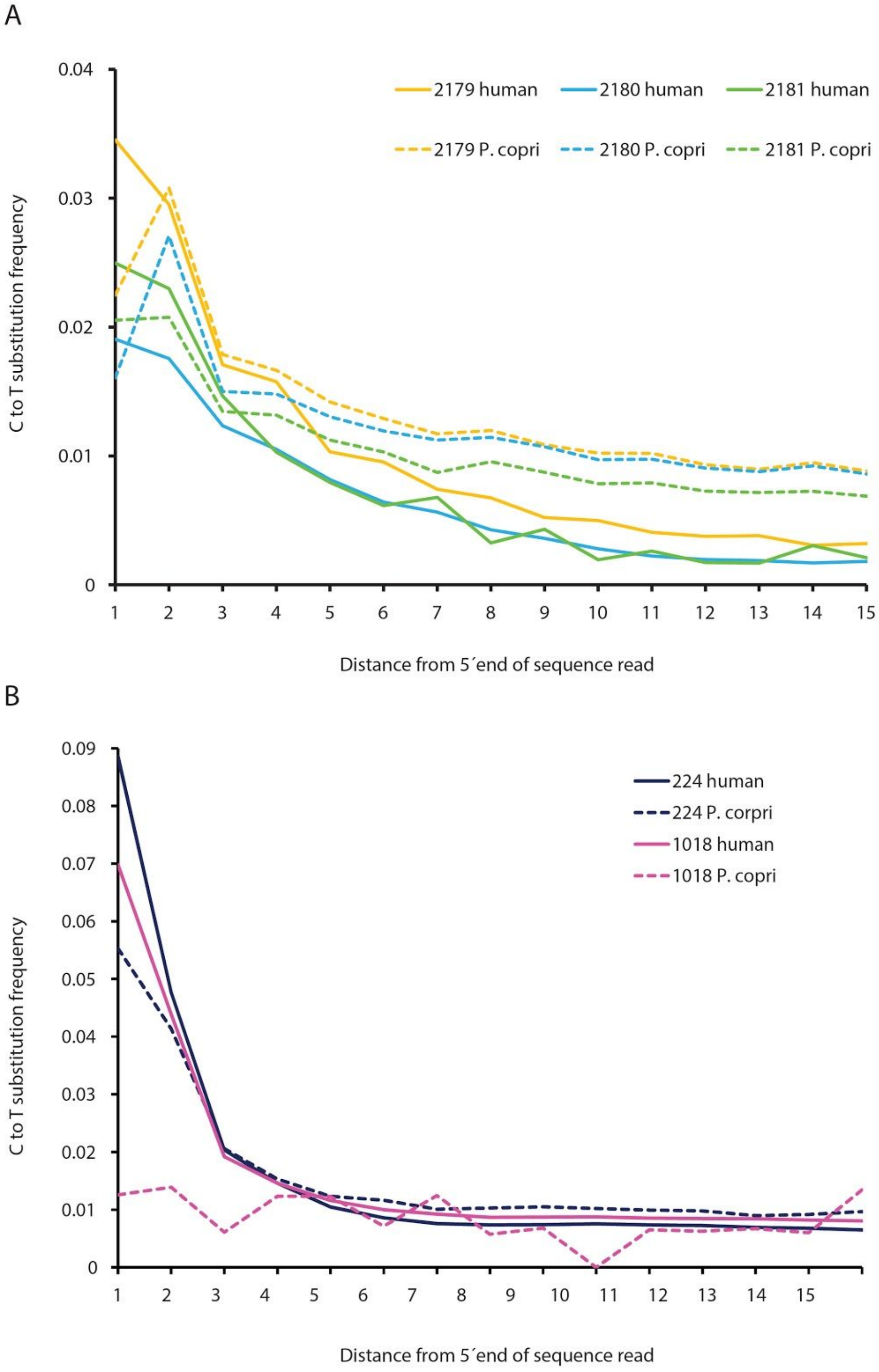


**Supplementary Figure 10**. Ancient DNA damage profiles. Cytosine to thymine substitution frequencies in the 5´end of the human and *P. copri* (dashed lines) sequence reads detected in the Mexican coprolite material **(A)** and in the Iceman samples **(B)**.

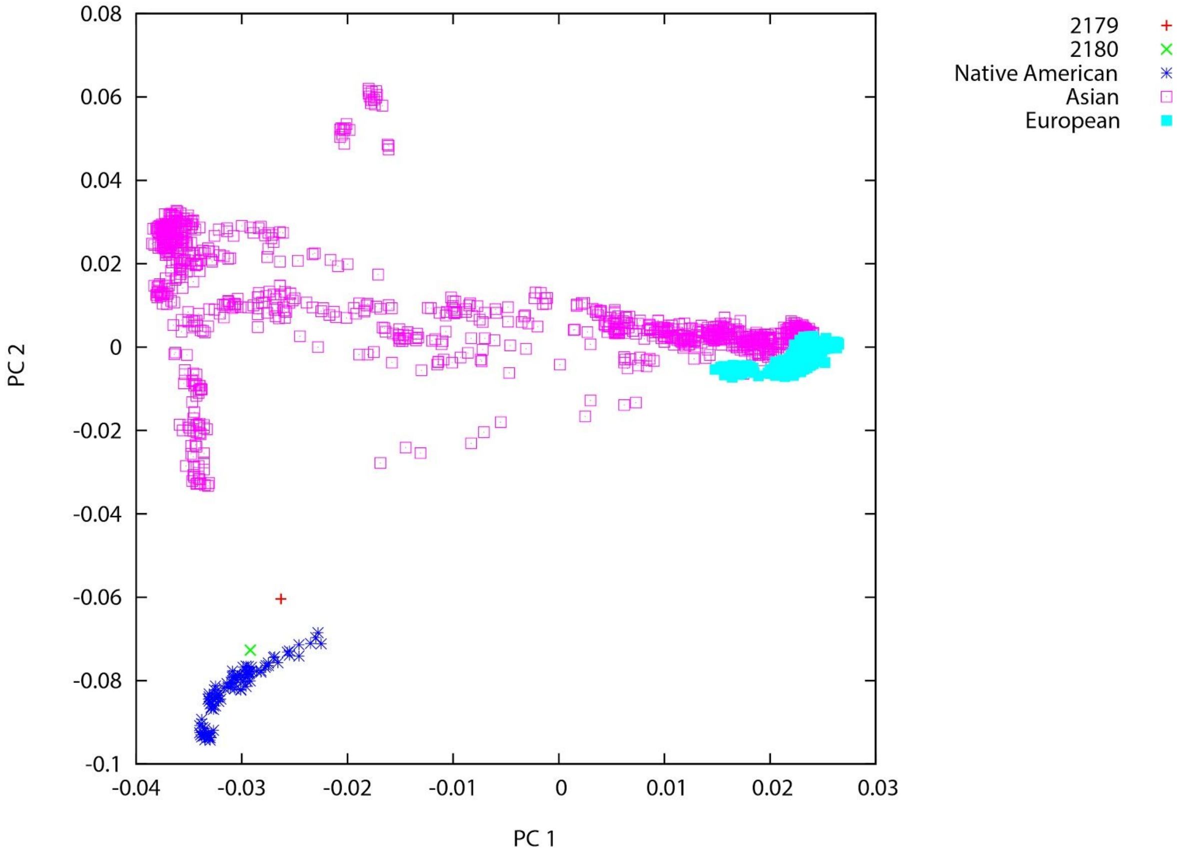


**Supplementary Figure 11**. PCA plot of two Mexican coprolite samples and selected modern European, Asian and Native American. Genome-wide ancient data was projected against a selected subset of the Affymetrix Human Origins populations.

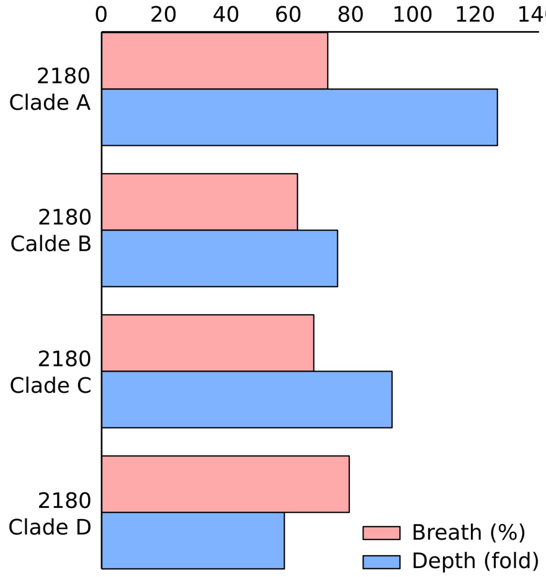


**Supplementary Figure S12**. Depth and breadth of metagenomic reads from sample 2180 mapped to four *P. copri* isolate genomes representing the four *P. copri* complex clades.
